# Selective roles of vertebrate PCF11 in premature and full-length transcript termination

**DOI:** 10.1101/491514

**Authors:** Kinga Kamieniarz-Gdula, Michal R. Gdula, Karin Panser, Takayuki Nojima, Joan Monks, Jacek R. Wisniewski, Joey Riepsaame, Neil Brockdorff, Andrea Pauli, Nick J. Proudfoot

## Abstract

The pervasive nature of RNA polymerase II (Pol II) transcription requires efficient termination. A key player in this process is the cleavage and polyadenylation (CPA) factor PCF11, which directly binds to the Pol II C-terminal domain and dismantles elongating Pol II from DNA *in vitro*. We demonstrate that PCF11-mediated termination is essential for vertebrate development. A range of genomic analyses, including: mNET-seq, 3’ mRNA-seq, chromatin RNA-seq and ChIP-seq, reveals that PCF11 enhances transcription termination and stimulates early polyadenylation genome-wide. PCF11 binds preferentially between closely spaced genes, where it prevents transcriptional interference and downstream gene silencing. Notably, PCF11 is sub-stoichiometric to the CPA complex. Low levels of PCF11 are maintained by an auto-regulatory mechanism involving premature termination of its own transcript, and are important for normal development. Both in human cell culture and during zebrafish development, PCF11 selectively attenuates the expression of other transcriptional regulators by premature CPA and termination.

## INTRODUCTION

RNA polymerase II (Pol II)-mediated transcription involves a cycle of initiation, elongation and termination. Transcription termination stops RNA synthesis through the release of Pol II and RNA from the DNA template. This process is crucial for correct gene expression for several reasons. First, termination punctuates the ends of transcription units and thereby releases the RNAs to fulfil their biological function. Second, it ensures Pol II availability for subsequent rounds of RNA synthesis. Termination also restricts the extent of non-coding transcription and prevents transcriptional interference between adjacent transcriptional units (Jensen et al., 2013; Porrua et al., 2016; Proudfoot, 2016).

Mechanistically, termination is coupled to the 3’ end processing of pre-mRNAs. Pol II becomes termination-competent after transcribing a polyadenylation signal (poly(A) signal, typically including the hexamer AAUAAA). This signal in the nascent RNA is recognized by the RNA 3’ processing machinery, which promotes RNA cleavage and polyadenylation 10-30 nucleotides downstream at the polyadenylation site (PAS) (reviewed in Proudfoot, 2011). Cleavage of the nascent transcript at the PAS is coupled to 5’->3’ degradation of the downstream RNA by XRN2, which eventually leads to termination (reviewed in Porrua et al., 2016; Proudfoot, 2016). In mammalian genomes Pol II typically continues transcribing for thousands of base pairs downstream of the PAS (Nojima et al., 2015; Schwalb et al., 2016). In summary, although the cleavage and polyadenylation (CPA) step occurs at defined locations (PAS), Pol II continues to transcribe the downstream sequences over a wide genomic window.

Most mammalian genes contain multiple alternative poly(A) signals. If more than one site within a transcript is able to support RNA cleavage, distinct RNA 3’ termini are generated, in a mechanism called alternative polyadenylation (APA) (Tian and Manley, 2017). APA within the coding region of genes can result in truncated polypeptides with diverse functions, as demonstrated for genes encoding calcitonin (Amara et al., 1984) and immunoglobulin heavy chain (Takagaki et al., 1996). Genome-wide studies have demonstrated that 70% of mammalian protein coding genes have alternative 3’ ends, which most frequently differ in their 3’ untranslated region (3’ UTR) of the mRNA. Alternative 3’ UTRs can confer different functionalities and stabilities to mRNAs depending on the presence of AU-rich elements and binding sites for miRNAs and RNA-binding proteins (reviewed in Tian and Manley, 2017).

3’ ends of mammalian mRNA are processed by a large cleavage and polyadenylation (CPA) complex, which includes the cleavage and polyadenylation specificity factor (CPSF), the cleavage stimulation factor (CstF), and cleavage factors I and II (CFIm and CFIIm), each consisting of multiple subunits (reviewed in Shi and Manley, 2015). CFIIm contains two proteins, PCF11 and CLP1, and - unlike other CPA factors - interacts only weakly and/or transiently with the complex (Shi et al., 2009). Most CPA proteins participate in defined steps, such as the cleavage reaction or recognition of specific RNA motifs. In contrast, PCF11 is critical not only for 3’ processing (Amrani et al., 1997; de Vries et al., 2000; Gross and Moore, 2001) but also for transcription termination (Zhang et al., 2005; Zhang and Gilmour, 2006; West and Proudfoot, 2008) and linking transcription with mRNA export (Hammell et al., 2002; Rougemaille et al., 2008; Johnson et al., 2009; Volanakis et al., 2017). In yeast the 3’ end processing and termination activities of PCF11 are provided by distinct PCF11 domains and can be functionally uncoupled (Sadowski et al., 2003). PCF11 is able to bind to the C-terminal domain (CTD) of the largest subunit of Pol II via its conserved CTD interaction domain (CID) (Barillà et al., 2001; Meinhart and Cramer, 2004; Kecman et al., 2018). The PCF11-CTD interaction dismantles elongation complexes *in vitro* (Zhang et al., 2005; Zhang and Gilmour, 2006) and is required for normal Pol II CTD serine 2 phosphorylation (S2ph) levels in yeast (Grzechnik et al., 2015), in line with CPA factor requirement for S2ph at the 3’ ends of human genes (Davidson et al., 2014). The CID of PCF11 also displays RNA binding activity, and a competition between RNA and CTD binding by the CID has been proposed to mediate Pol II disengagement (Zhang et al., 2005; Hollingworth et al., 2006). A second RNA-binding domain is present in the C-terminal part of the protein (Schäfer et al., 2018).

Although PCF11 is a key factor acting at the intersection of several nuclear processes, it has mainly been studied in yeast, with little knowledge of its function in vertebrates. However, three independent pan-cancer screens for cancer driver mutations have recently identified recurrent mutations in *PCF11*, in particular within the promoter region (Hornshøj et al., 2018; Kuipers et al., 2017; Rheinbay et al., 2017). Moreover, PCF11 expression levels are predictive of clinical outcomes of neuroblastoma patients (Ogorodnikov et al., 2018), suggesting that PCF11 has relevance to human pathology. We address here the genome-wide role of PCF11 in vertebrate gene expression.

## RESULTS

### PCF11 enhances transcription termination and CPA genome-wide

We first tested whether PCF11 acts as a general transcription termination factor in human cells. PCF11 was depleted using a pool of four siRNAs optimized for knock-down duration and siRNA dosage (Figure S1A-B), to limit indirect effects. To assess Pol II binding we employed ChIP-seq for total Pol II (using the N20 antibody). Transcriptional output was also measured by analysis of chromatin-bound RNA (chrRNA), which is enriched for nascent transcripts. We further employed mNET-seq (Nojima et al., 2015) to assay nascent transcripts associated with the threonine 4 phosphorylated (T4ph) Pol II CTD, which is specific to the termination region (Heidemann et al., 2013; Schlackow et al., 2017).

PCF11 depletion led to transcriptional read-through beyond usual end sites (Figure 1A-C, S1C-D). While many genes show Pol II accumulation downstream the PAS, PCF11 depletion shifted and decreased this Pol II pausing (Figure 1A-C). Transcript read-through and altered Pol II pausing are both indicative of defective termination. While all assays consistently showed delayed termination in PCF11 depleted conditions, T4ph mNET-seq provided the most specific detection of transcriptional termination. PCF11-depletion induced termination delay was wide-spread, but only resulted in a shift in the termination window rather than a complete failure to terminate. Downregulation of other human CPA and termination factors in previous studies led to a similar termination shift, suggesting the existence of uncharacterized failsafe termination mechanisms in mammals (Fong et al., 2015; Nojima et al., 2015; Schlackow et al., 2017).

**Figure 1.**
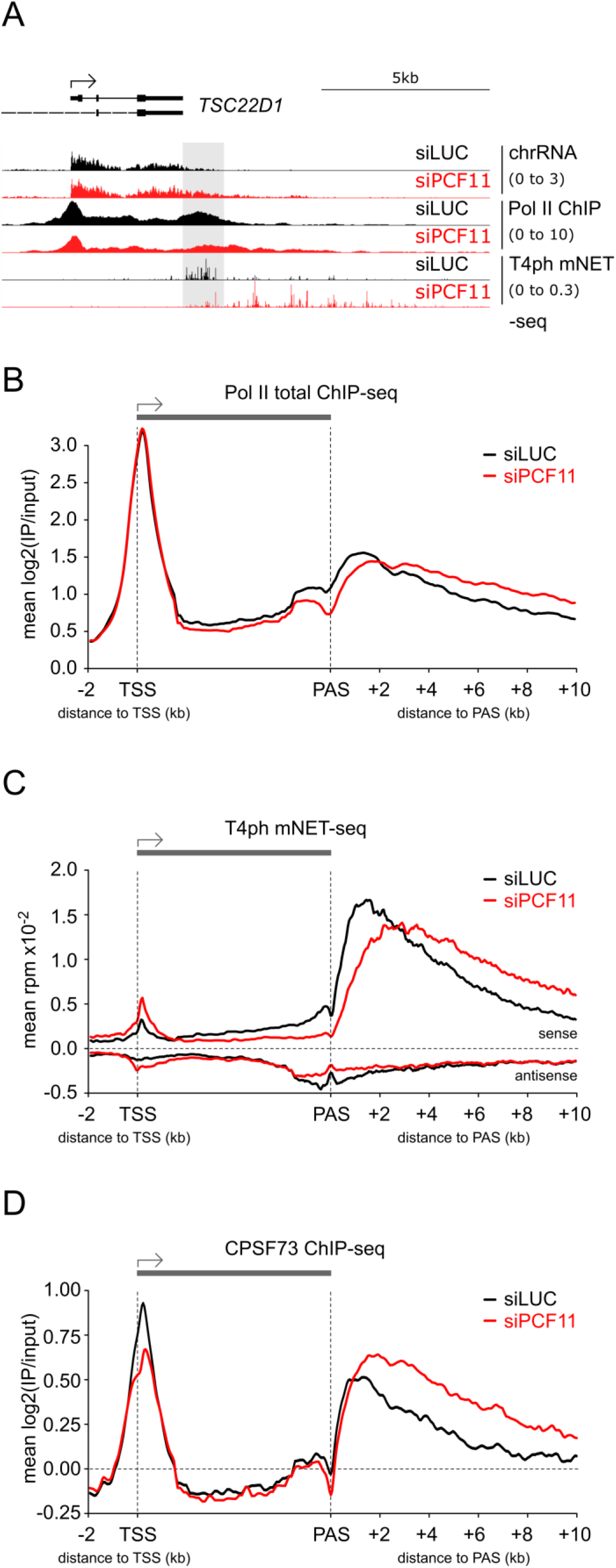
PCF11 enhances transcription termination genome-wide. (A) Genomic profile of *TSC22D1*. Grey shading highlights the termination window in control cells (siLUC, black). PCF11 depletion (siPCF11) is depicted in red. For chrRNA-seq and mNET-seq only sense strand is shown, Pol II ChIP-seq is not strand specific. In all profiles: numbers in brackets indicate the viewing range (rpm).
(B-D) Metagene analysis of total Pol II ChIP-seq (B), T4ph mNET-seq (C) and CPSF73 ChIP-seq (D) in cells ± siPCF11 on transcribed protein-coding genes >5 kb long which PAS is isolated >6 kb downstream from the nearest gene on the same strand (n=8389).

The observed termination delay upon PCF11 depletion could be due to a concomitant loss of CPA complex association with termination regions. To test this hypothesis we performed ChIP-seq for CPSF73 (CPSF3), which is the CPA subunit responsible for pre-mRNA 3’ cleavage. Contrary to expectations, PCF11 depletion resulted in increased and 3’ extended CPSF73 signal at gene 3’ ends (Figure 1D, S1E), consistent with prolonged CPSF73 binding to chromatin. Therefore, PCF11 may not be necessary for CPA complex binding to PAS-proximal regions, but instead appears to increase CPA efficiency. In conclusion, our data indicate that human PCF11 enhances genome-wide CPA and transcription termination.

### PCF11-mediated termination enhancement occurs independently of APA

PCF11 not only affects CPA and termination, but also regulates APA (Li et al., 2015). The read-through we observed upon PCF11 depletion could either be a *bona fide* termination defect, or due to a shift towards distal PAS usage. To distinguish these possibilities, we determined active PAS usage by sequencing the 3’ ends of nuclear polyadenylated RNAs (3’ mRNA-seq) from control and PCF11 depleted cells. A set of 11947 protein-coding and non-coding genes, with PAS at least 6kb away from the downstream gene on the same strand was selected. PCF11 depletion caused a shift to distal PAS usage for 2072 genes (17%), while only 333 (2.8%) revealed a proximal shift, indicating that PCF11 favours proximal PAS usage in human cells (Figure 2A-C, S2A-B). This effect was more pronounced for protein-coding (22% distal and 3% proximal shifts), than non-coding genes, of which only 4.4% showed any PAS shift (Figure S2A). Genes undergoing APA changes upon PCF11 knock-down had overall higher expression levels than genes where no shift occurred (Figure S2C). Analysis of the 3’ mRNA-seq data furthermore revealed global gene down-regulation upon PCF11 depletion (Figure S2D). The majority of genes with significantly altered expression upon PCF11 knock-down showed no shift in APA usage (Figure S2D, red dots), which suggests that differential PAS usage is not the major cause of siPCF11-induced gene deregulation. An important outcome from our APA analysis is that termination loss following PCF11 depletion often occurs without a distal APA shift, and that the termination window shifts downstream in all four categories of PAS usage (Figure 2D-E, S2E). Therefore we conclude that read-through transcription upon PCF11 depletion is a hallmark of delayed termination, and not generally due to differential PAS selection.

**Figure 2.**
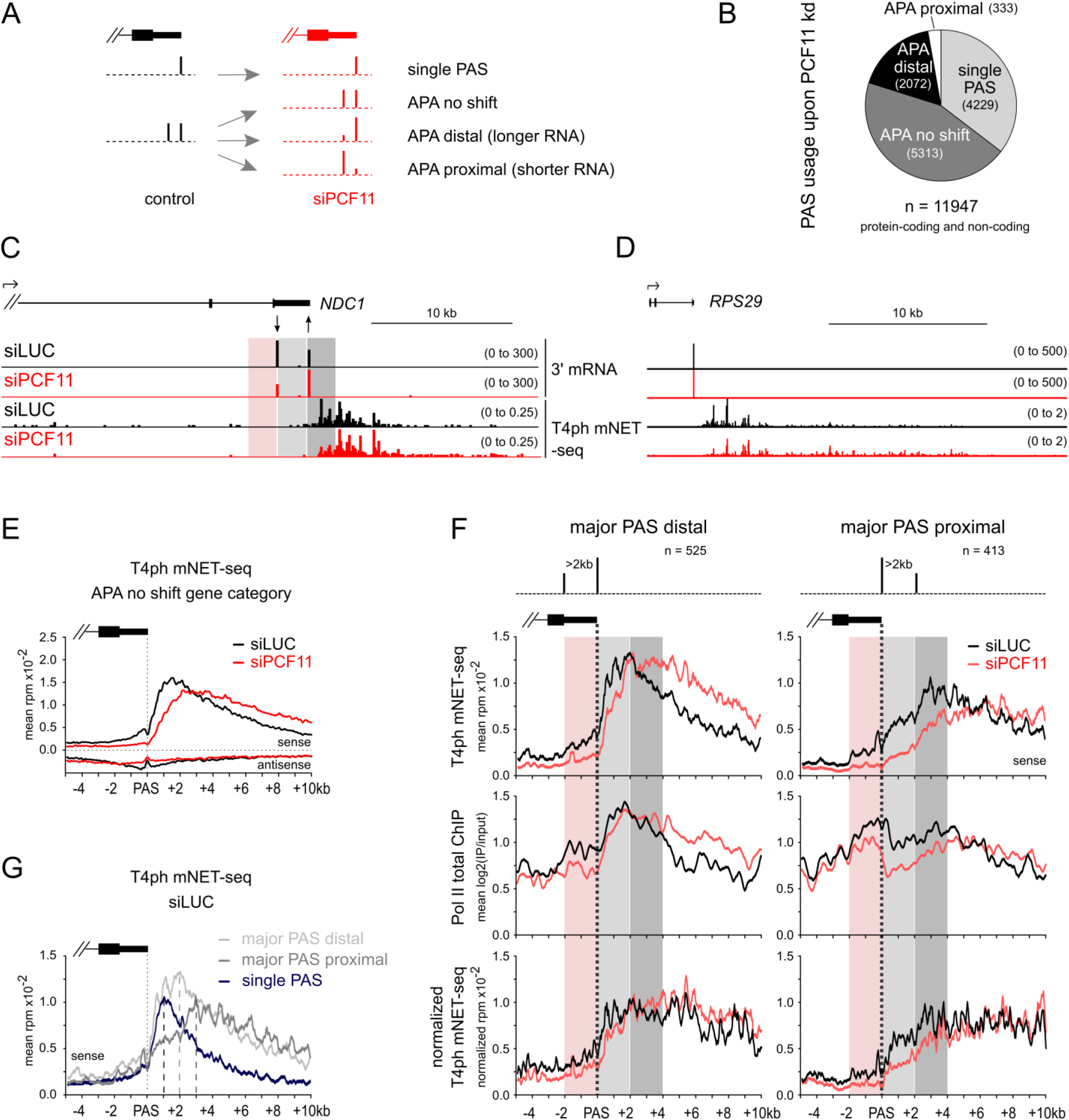
PCF11-mediated termination enhancement occurs independently of PAS selection. (A) Schematic of gene categorization based on APA changes ± siPCF11.
(B) Pie chart of PAS usage in cells depleted of PCF11 based on DEXseq analysis (p-adjusted < 0.05). (C-D) Genomic profiles of *NDC1* and *RPS29*. PASs indicated by arrows show significant APA in 4 repeats of 3’mRNA-seq ± PCF11 (DEXseq p-adjusted < 0.05), Figure S2B.
(C and F) Red shading highlights the region 2kb upstream of the major PAS, light grey 2kb downstream and dark grey 2-4kb downstream of the major PAS.
(E-G) Meta-gene profiles of T4ph mNET-seq signal – 5kb to +10 kb around the major PAS on protein-coding genes. Vertical dotted line highlights the position of the major PAS.
(E) Multiple PASs-containing genes without significant change in PAS usage (APA no shift); meta-profiles for other APA gene categories shown in Figure S2E.
(F) Genes with two strongest PASs of comparable signal and separated by > 2kb (n = 938) were divided into two sets: those with a major distal (left panels) or proximal (right panels) PAS. Schemes of the relative positioning of the two strongest PASs are shown on top. Top panels: T4ph mNET-seq, middle panels: Pol II total ChIP-seq, bottom panel: T4ph mNET-seq normalized to Pol II total.
(G) Comparison of T4ph mNET-seq profiles in control cells (siLUC) for the indicated gene categories. Vertical dashed lines highlight the corresponding T4ph mNET-seq signal maxima.

Intriguingly, we observed numerous genes with well-separated PASs but no T4ph mNET-seq signal associated with the proximal PAS (Figure 2C, S2F-G, light grey shading). To determine whether this is a general trend, we selected a subset of 938 protein-coding APA genes where the two strongest PASs differed no more than 2-fold, and were separated by at least 2 kb. If the strongest of the two PASs was proximal, the gene was classified as “major PAS proximal”, otherwise as “major PAS distal” (Figure 2F). Genes where the major PAS is distal showed a sharp T4ph mNET-seq signal increase immediately after the PAS, coinciding with higher Pol II density in the same region. Conversely, Pol II density decreased downstream of genes where the major PAS is proximal, and T4ph mNET-seq signal increased more gradually. PCF11 depletion caused delayed termination for both gene categories (Figure 2F). In control cells, the highest T4ph mNET-seq signal occurred on average 2 kb downstream of the PAS for genes with a major distal PAS and 3 kb for genes with a major proximal PAS, as compared to 1 kb for genes with only 1 PAS (Figure 2G). When normalized to Pol II density, T4ph mNET-seq signal reached a plateau within 0.5 kb from the PAS for major PAS distal genes, and after 2.5 kb for major PAS proximal genes (Figure 2F, bottom panels). We conclude that T4ph mNET-seq is more closely associated with distal PASs. These data support the previous suggestion that CPA and termination might be uncoupled, and are also consistent with the possibility of PAS choice occurring in favour of proximal PAS even after the distal PAS has been transcribed (Zhu et al., 2018). Further work involving long-read sequencing is needed to verify whether transcription generally terminates downstream of the distal PAS.

Overall, our genomic analyses reveal that PCF11 plays a specific role in enhancing transcription termination independently of APA selection.

### Genomic binding pattern of PCF11

To determine at which point in the transcription cycle PCF11 binds to Pol II, we analyzed the genomic binding profile of PCF11 by ChIP-seq. Two antibodies specifically enrich for PCF11 in IPs (Figure 3A, S3A-B): one targeting a C-terminal epitope (PCF11-Ct), the other one an internal epitope (PCF11-Int). Both gave similar ChIP-seq profiles (Figure S3C-E). Consequently, merged signals from both antibodies are shown, labelled PCF11-(Int+Ct) (Figure 3B). We only considered regions significantly above background for both antibodies (1% FDR) as PCF11 enriched (Figure 3C).

**Figure 3.**
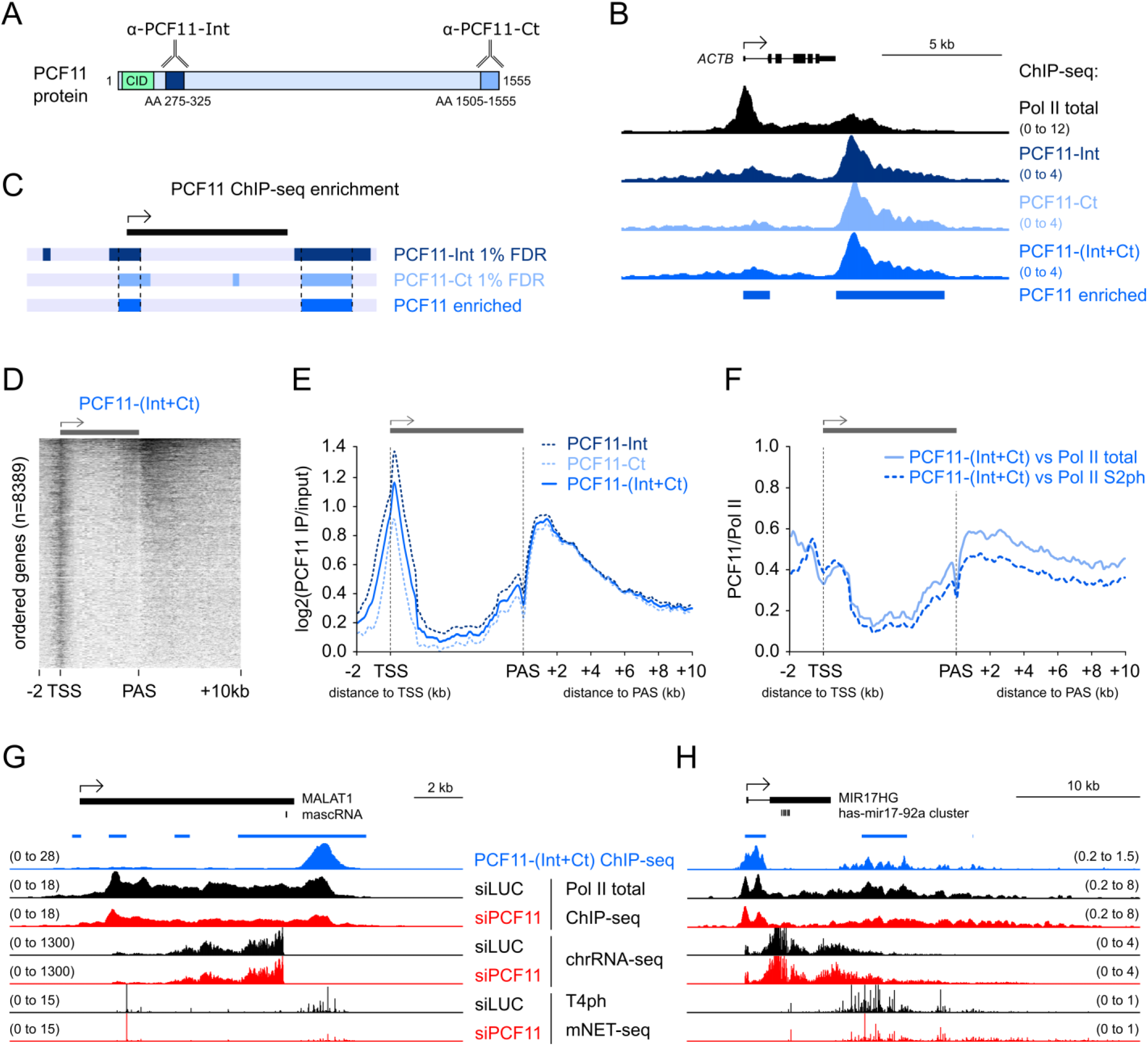
Genomic pattern of PCF11 binding. (A) Position of epitopes recognized by PCF11 antibodies; α-PCF11-Int binds an Internal peptide, and α-PCF11-Ct the C-terminus. Amino acid (AA) numbering corresponds to the main human PCF11 isoform NP_056969.2.
(B) Genomic profile of PCF11 binding to the *ACTB* gene. PCF11-(Int+Ct) corresponds to the merged antibodies profile. Blue bars below PCF11 ChIP-seq signal indicate PCF11 enrichment.
(C) Regions with ChIP-seq signal significantly above background for both PCF11 antibodies were considered PCF11-enriched.
(D) Heatmap of PCF11-(Int+Ct) ChIP-seq signal (log2IP/input) across protein-coding genes ranked from highest to lowest PCF11 signal.
(E-F) Meta-gene analysis of PCF11 binding on protein-coding genes. Plotted are average log2 ratios of PCF11 ChIP-seq signal relative to chromatin input (E) and relative to Pol II (F). PCF11 and Pol II signals were calculated as log2(IP/input).
(G-H) Genomic profiles of *MALAT1* and *MIR17HG* showing PCF11 binding and the transcriptional effect of PCF11 depletion.

Most prominent PCF11 binding occurred at 3’ end of genes, downstream of the PAS (Figures 3B-D and S3D,E), fitting with a role for PCF11 in termination. However, TSS-proximal enrichment was more frequent (Figure 3D, S3F), leading to a global binding profile at both gene ends (Figure 3E). Consistent with PCF11 binding to the TSSs, upon PCF11 depletion T4ph mNET-seq signal was not only altered at the 3’ end but additionally increased in both sense and antisense direction at the TSS (Figure 1C). Non-coding TSS-associated transcription units, unlike protein-coding genes, are uniformly marked by Pol II T4ph (Schlackow et al., 2017), therefore the observed increased levels of TSS-associated T4ph mNET-seq are indicative of increased transcription upon PCF11 depletion. This is in line with the previously published role of termination factors in restricting non-productive RNA synthesis at the TSS (Nojima et al., 2015). The pattern of PCF11 binding across genes relative to Pol II (Figure 3F, S3F) suggests that PCF11 doesn’t consistently travel with elongating Pol II from the promoter to the PAS, although it could transiently interact with Pol II across the gene body.

Interestingly, while CPA-dependent protein-coding genes were the major target of PCF11 binding, high levels of PCF11 were also detected on transcription units using alternative 3’ processing mechanisms (Figures S3G, 3G-H). For example, the 3’ end of RNase P-processed *MALAT1* gene is one of the top loci enriched for PCF11. PCF11 down-regulation did not affect transcription of this gene, but decreased Pol II and T4ph mNET-seq signals (Figure 3G). Most other non-canonical PCF11 targets also showed no read-through transcription upon siPCF11, with the exception of microprocessor-dependent lncRNA microRNA host genes (Dhir et al., 2015) like *MIR17HG* (Figure 3H). PCF11 binding to CPA-independent genes could point to non-canonical functions for PCF11/CPA factors on these transcripts, or a hybrid mechanism where alternative 3’ processing pathways are used in parallel.

### Closely spaced genes are PCF11-dependent

Although PCF11 binding was detectable on both coding and non-coding transcription units, only 54% of tested polyadenylated protein-coding genes had significant PCF11 enrichment (4516/8389), of which 47% (2140) showed enrichment in the 3’ end region. PCF11-enriched genes were globally more highly expressed compared to non-enriched genes (Figure S4A). However, PCF11 enrichment occurred at some silent loci, and vice versa, some highly expressed genes had no enrichment. For example, PCF11 was enriched at the *ZNF786* but not the *PDIA4* termination region (Figure 4A), although the latter is transcriptionally more active (chrRNA-seq) and produces >200 fold more polyadenylated nuclear mRNA (3’ mRNA-seq). *PDIA4* showed a high exonic/intronic chrRNA ratio, implying fast splicing, and had no Pol II accumulation downstream of the PAS. We hypothesize that rapidly processed genes may be bound by PCF11 only transiently.

**Figure 4.**
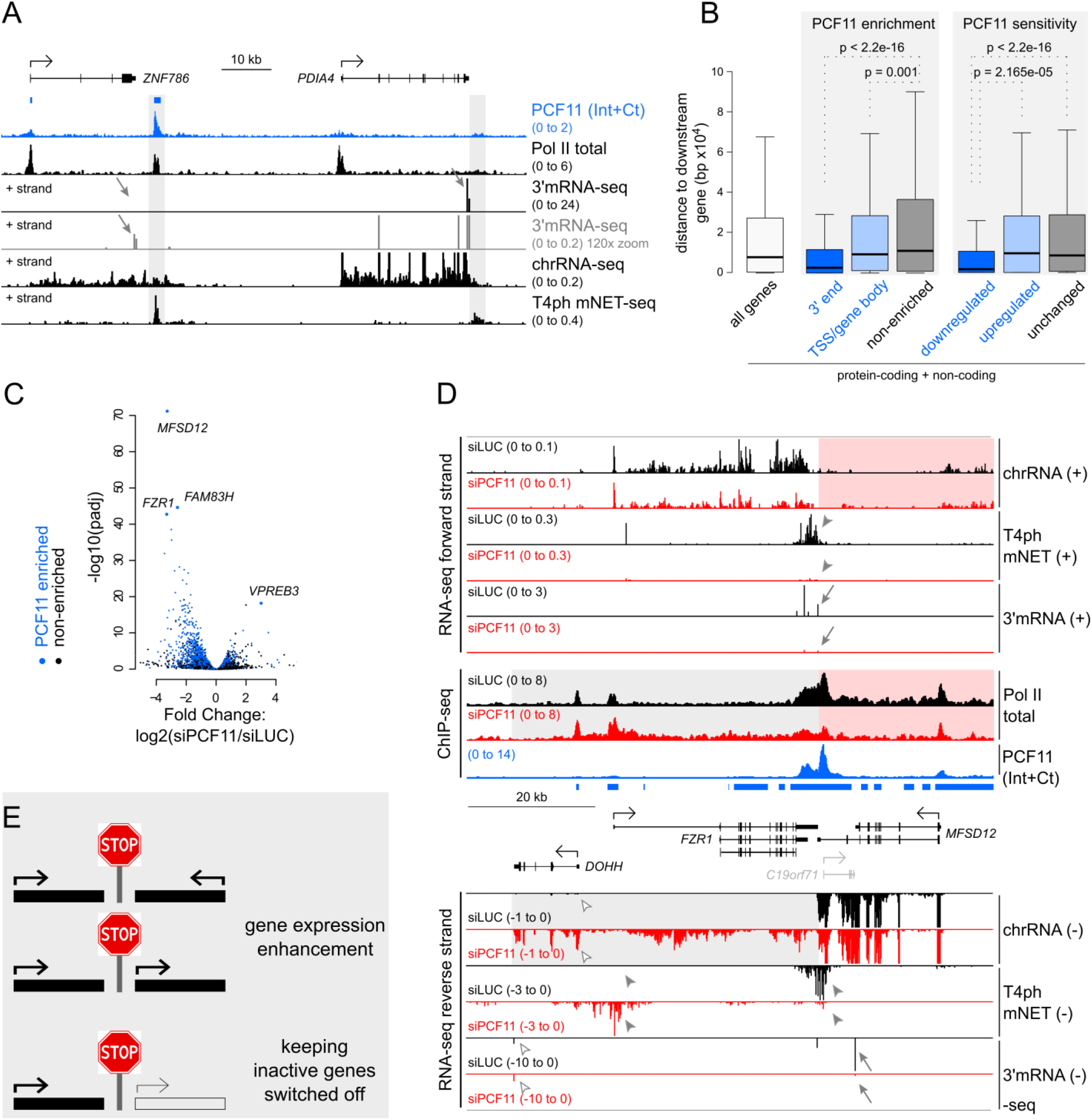
Closely spaced genes are PCF11-dependent. (A) Genomic profile of *ZNF786/PDIA4*. Blue bars on top indicate PCF11 enrichment. The termination regions are shaded. 3’ mRNA-seq data is shown at two viewing ranges.
(B) Boxplot showing the distances between genes’ PASs to their nearest gene downstream. Statistical significance was determined using Mann-Whitney test. Here and in all boxplot figures, the thick horizontal line marks to the median, and the upper and lower limits of the box the 1^st^ and 3^rd^ quartile.
(C) Volcano plot showing differential expression between the siLUC and siPCF11 conditions. Blue dots correspond to PCF11 enriched genes, black dots to non-enriched genes. The four most significantly differentially regulated genes are indicated.
(D) Genomic profile of the *FZR1/MFSD12* locus. Data from the + strand and non-strand specific ChIP data are shown above the locus, data from the – strand below it. Grey shading highlights the read-through of *MFSD12*, red shading lack of detectable read-through from *FZR1*. *FZR1* is less active (10x zoomed in viewing range). Blue bars: PCF11 enriched regions; arrows: gene downregulation measured by 3’ mRNA-seq; filled arrowheads: alterations in T4ph mNET-seq signal; empty arrowheads: upregulation of *DOHH* gene due to read-through transcription from *MFSD12*.
(E) Model of PCF11s role in enhancing gene expression of closely spaced genes and isolating inactive genes from upstream tandem active genes.

Visual inspection of PCF11 ChIP-seq data revealed high PCF11 levels between closely spaced genes (Figure S3D-E). Accordingly, genes with 3’ PCF11-enrichment showed 4-fold smaller median spacing compared to non-enriched genes (Figure 4B, S4B). We hypothesize that PCF11 enrichment between closely spaced transcription units prevents transcriptional interference between adjacent genes. Supporting this view, genes significantly downregulated upon siPCF11 treatment were 5-fold more closely spaced than PCF11-insensitive genes (Figure 4B).

Two of the three most downregulated genes, *MFSD12* and *FZR1*, are convergent neighbours (Figure 4C-D). PCF11 showed enrichment over a large part of the locus, especially pronounced in between them (Figure 4D). Interestingly, while PCF11 depletion abrogated the formation of polyadenylated mRNA products for both genes, transcription, as measured by chrRNA-seq and Pol II ChIP-seq levels, was only mildly decreased. At the same time, strong read-through transcription of the more highly transcribed *MFSD12* gene was evident (grey shading). Furthermore, T4ph mNET-seq signal was abrogated for *FZR1* and shifted >20kb for *MFSD12*. This is in contrast with isolated genes, where termination is typically shifted only mildly (compare with Figures 1A, 2C-D). The similar chrRNA levels in the *MFSD12*/*FZR1* locus upon depletion of PCF11 suggest that loss of mRNA is not due to transcriptional inhibition, but rather a result of failure of 3’ processing accompanying the severe termination defect. Extensive transcriptional interference is not limited to convergent genes, but can also affect the downstream gene of transcription units oriented in tandem, such as *TIMM13* (Figure S4C). Notably, the top two deregulated genes by siPCF11, *MFSD12* and *FAM83H*, showed loss of mRNA in the absence of detectable read-through transcription from the lower or not expressed close-by gene – *FZR1* and *MAPK15* (Figure 4D and S4D, red shading). We speculate that the presence of a close-by downstream gene, even if it is inactive, doesn’t allow for fail-safe termination mechanisms to compensate for PCF11 downregulation. This leads to a failure in both 3’ end processing and termination.

Even though PCF11 depletion causes a global downregulation of genes, some genes were upregulated (Figure 4C, log2(siPCF11/siLUC) > 0). Visual inspection of these genes (e.g. top upregulated *VPREB3*) revealed that upon PCF11 depletion many were invaded by read-through transcription from an upstream tandem gene (Figure S4E, see also *DOHH* in Figure 4D). These events likely result in unproductive, fused transcripts, but could in some cases (e.g. poised genes) also activate independent transcription of the downstream gene by altering the chromatin environment.

Overall, we predict that PCF11-mediated gene punctuation is essential to promote efficient gene expression of closely spaced active genes. Additionally, it isolates inactive genes from active upstream genes (Figure 4E).

### PCF11 is substoichiometric to the CPA complex

The selective pattern of PCF11 enrichment on genes (Figure 4) prompted us to assess PCF11 protein levels relative to Pol II and other components of the 3’ processing machinery, taking advantage of global quantitative proteomics. Surprisingly, we found that PCF11 is substoichiometric to other CPA complex subunits across different human cells and tissues, measuring on average 10-20 fold less molecules per cell than other CPA and Pol II subunits (Figure 5A and S4F). PCF11 is also the least abundant human CID-containing protein (Figure S4G). Consistent with the proteomics data, *PCF11* mRNA levels were among the least abundant and most unstable CPA mRNAs, as measured by BRIC-seq (Figure 5B) (Tani et al., 2012). The unusually low protein and mRNA level of PCF11 implies possible transcriptional regulation. Strikingly, *PCF11* has an evolutionarily conserved first intron, which harbours highly conserved tandem AATAAA hexamers flanked by upstream and downstream elements that aid CPA factor recruitment (Figure 5C and S5A). These sequences together form a highly active PAS, referred to as PAS1 (Figure 5C). The short PCF11 isoform resulting from PAS1 usage encodes a short ORF with a C-terminal extension. However, this polypeptide was not detected by mass spectrometry analysis in HeLa, U2OS and HEK 293 cell lines.

**Figure 5.**
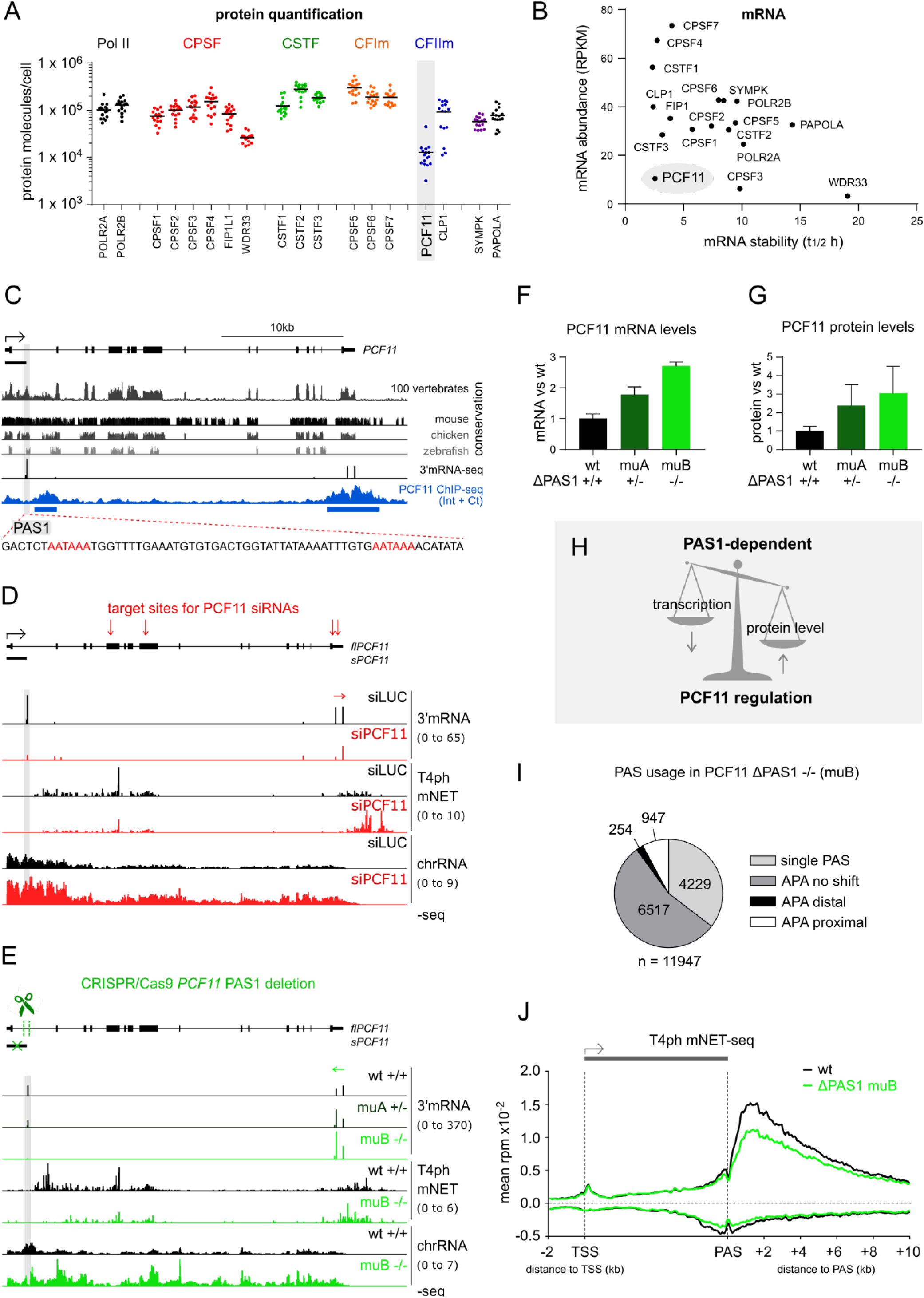
PCF11 is substoichiometric to the CPA complex and autoregulates its levels by premature CPA and termination. (A) Scatter plot of protein molecules per cell of Pol II subunits and CPA subcomplexes from global quantitative proteomics (Wiśniewski et al., 2015). Each dot represents a colorectal adenoma biopsy from a different patient (n=16). Horizontal lines corresponds to the mean. For data from other biopsies and HeLa cell culture see Figure S4F.
(B) Scatter plot of mRNA abundance versus stability of the same factors as (A) in HeLa cells based on BRIC-seq (Tani et al., 2012).
(C) Genomic profile of *PCF11* showing its evolutionary conservation in 100 vertebrates (top track) and in individual species (middle tracks). Actively used PASs measured by 3’mRNA-seq, PCF11 ChIP-seq signal and PCF11 enrichment (blue boxes) are shown for human cells. Viewing range was auto-scaled to data. Grey shading highlights the conserved PAS in the first intron. Underlying DNA sequence below showing tandem AATAAA hexamers in red.
(D) PCF11 depletion affects transcription of its own gene. Top: schematic of *PCF11* indicating locations of siRNA target sites (vertical red arrows). Tracks: comparison of 3’mRNA-seq, T4ph mNET-seq and chrRNA-seq ±PCF11. Horizontal arrows (D and E) show the direction of APA.
(E) *PCF11* PAS1 deletion affects transcription of the *PCF11* gene. Top: schematic of *PCF11* indicating a 285 bp CRIPR/Cas9-mediated deletion in the 3 kb first intron, removing the tandem poly(A) signals. Tracks are as in (D) for wild-type (wt) cells and CRISPR/Cas9 clones with a partial deletion (muA +/−) and full deletion (muB −/−) of PAS1. Further clones are shown in Figure S5E.
(F) Quantification of full length *PCF11* mRNA levels in wt and PAS1 deletion clones based on the 3’mRNA-seq count in the *PCF11* 3’UTR (error bars correspond to SD, n=3).
(G) Quantification of PCF11 WB signal in wt and PAS1 deletion clones. Error bars correspond to SD from three independent WB experiments; all samples were loaded in two dilutions in each experiment (n=6). Representative WB and additional deletion clones are shown in Figure S5F.
(H) Model: PCF11 protein levels modulate transcription of the *PCF11* gene in a PAS1-dependent manner, allowing for PCF11 autoregulation.
(I) Pie chart of genome-wide PAS usage and APA occurrence in ΔPAS1 clone muB vs wt cells (n=11947 genes, compare with Figure 2B).
(J) Meta-gene analysis of T4ph mNET-seq signal in wt cells and ΔPAS1 clone muB.

Interestingly, PCF11 was not only enriched at the 3’ end of its own gene, but also downstream of PAS1 (Figure 5C, blue track), suggesting that its low expression could be due to autoregulation by APA and premature termination.

### PCF11 is autoregulated by premature CPA and termination

Autoregulation of PCF11 by premature termination predicts that downregulation of PCF11 should lead to a decrease in PAS1 usage. We therefore knocked down PCF11 by siRNAs that specifically targeted full-length *PCF11* RNA (*flPCF11*) but not the short *sPCF11* isoform (Figure 5D, red vertical arrows). Notably, PAS1 usage and *sPCF11* levels decreased 5-fold upon PCF11 depletion, two times more compared to the directly targeted *flPCF11* (Figure 5D and S5B). Accordingly, T4ph mNET-seq signal downstream of PAS1 decreased, and instead increased at the 3’ end of *PCF11*. Also chrRNA signal increased across the whole gene (Figure 5D). This suggests that PAS1 usage depends on PCF11 levels and that PAS1-linked premature termination regulates *flPCF11* transcription. Interestingly, PAS1 appears particularly sensitive to PCF11 levels, compared to other 3’ processing factors. Re-analysis of data from a published mouse database (Li et al., 2015) reveals that depletion of mCFI-68, mPABPC1 and mPABPN1 increased PAS1 usage in nuclear RNA, whereas mFip1 depletion caused a smaller reduction compared to mPcf11 (Figure S5C).

To directly demonstrate the autoregulatory role of *PCF11* PAS1, we specifically deleted PAS1 including its flanking sequences (285bp) from the ~3kb intron by CRISPR/Cas9 (Figures 5E and S5D). Since we obtained only one full *PCF11ΔPAS1* mutant clone (muB) out of ~100 single-cell colonies tested, we also included in our analysis three partial deletion mutant clones (muA, muC and muD; Figure S5D). Consistent with a negative regulatory role of PAS1 on PCF11 expression, all four mutant clones displayed increased *flPCF11* mRNA levels as measured by 3’ mRNA-seq (Figure 5E-F, S5E) and an increase in PCF11 protein levels (Figure 5G and S5F). Additionally, T4ph mNET-seq of clone muB showed that PAS1 deletion caused a downregulation of *PCF11* intragenic T4ph mNET-seq signal with a concomitant 3’ end increase (Figure 5E). Moreover, chrRNA signal increased across the *PCF11* gene in muB cells (Figure 5E).

In conclusion, PAS1-linked APA and premature termination enable the cell to balance *PCF11* transcription to maintain stable and relatively low levels of PCF11 protein expression (Figure 5H).

Since the deletion of *PCF11* PAS1 induces PCF11 overexpression, we examined the mutant clones with respect to their genome-wide effect on APA and transcription termination. PCF11 overexpressing cells show a preference for proximal APA usage (Figure 5I and S5G) and a smaller window of T4ph mNET-seq signal (Figure 5J). These features are consistent with early CPA and termination caused by the increased levels of PCF11. PCF11-dosage-dependent effects on APA and termination are exemplified by *PCF11* itself: 3’UTR PAS usage and T4ph mNET-seq signal both shifted distally upon PCF11 reduction (Figure 5D) and proximally upon PCF11 levels increase (Figure 5E). T4ph mNET-seq signal was specifically downregulated at 3’ ends of genes in muB cells (Figure 5J), however neither PCF11 depletion nor upregulation affected global T4ph levels (Figure S5I). The decrease in T4ph mNET-seq signal might possibly be a result of more efficient CPA due to high PCF11 levels. PCF11 depletion had a stronger effect than PCF11 upregulation – with more APA distal shifts vs proximal shifts (Figure 2B vs 5I) and strong downregulation of gene expression vs. mild upregulation (Figure 4C vs S5H). In conclusion, PCF11 depletion and overexpression show opposite genome-wide effects on APA and termination, supporting the view that they are directly controlled by PCF11.

### PCF11 is essential and undergoes PAS1-dependent autoregulation during vertebrate development

To extend our findings to a physiological context, we analysed PAS1 usage in human tissue. Ranking of 22 tissues according to *PCF11* mRNA levels revealed wide-spread usage of PAS1, with the notable exception of 4 tissues with low *PCF11* expression (Fig. S5J). We therefore tested the importance of PCF11 and its PAS1-dependent regulation *in vivo*. We chose zebrafish as model organism as the *PCF11* PAS1 is highly conserved (Figure 5C) and actively used in this species (Figure S6A). To assess PCF11 function in vertebrate development, we inactivated zebrafish *pcf11* (*zPCF11*) by generating a CRISPR/Cas9-mediated frame-shift mutation (68 bp insertion) in the first coding exon of *zPCF11* (*zPCF11^null^*, Figure 6A, S6B-C). While embryos and adult fish heterozygous for the mutation (*zPCF11^null^*+/−) were indistinguishable from wild-types (+/+), incrosses of *zPCF11^null^*+/− fish resulted in ~25% of *zPCF11^null^*−/− embryos with severe brain and central nervous system necrosis by 20 hours post fertilization (hpf) (Figure S6D and 6B), which resulted in death within four days. The fully penetrant brain necrosis could be rescued by *zPCF11* mRNA injection (150 pg) (Figure 6B-C), confirming that the defects are due to loss of zPCF11. The initially normal development of *zPCF11^null^*−/− embryos is likely due to the maternal deposition of zPCF11 in the egg (Figure S6E), which supports normal development for the first hours. Together, our analysis provides the first direct evidence that PCF11 is essential for vertebrate development.

**Figure 6.**
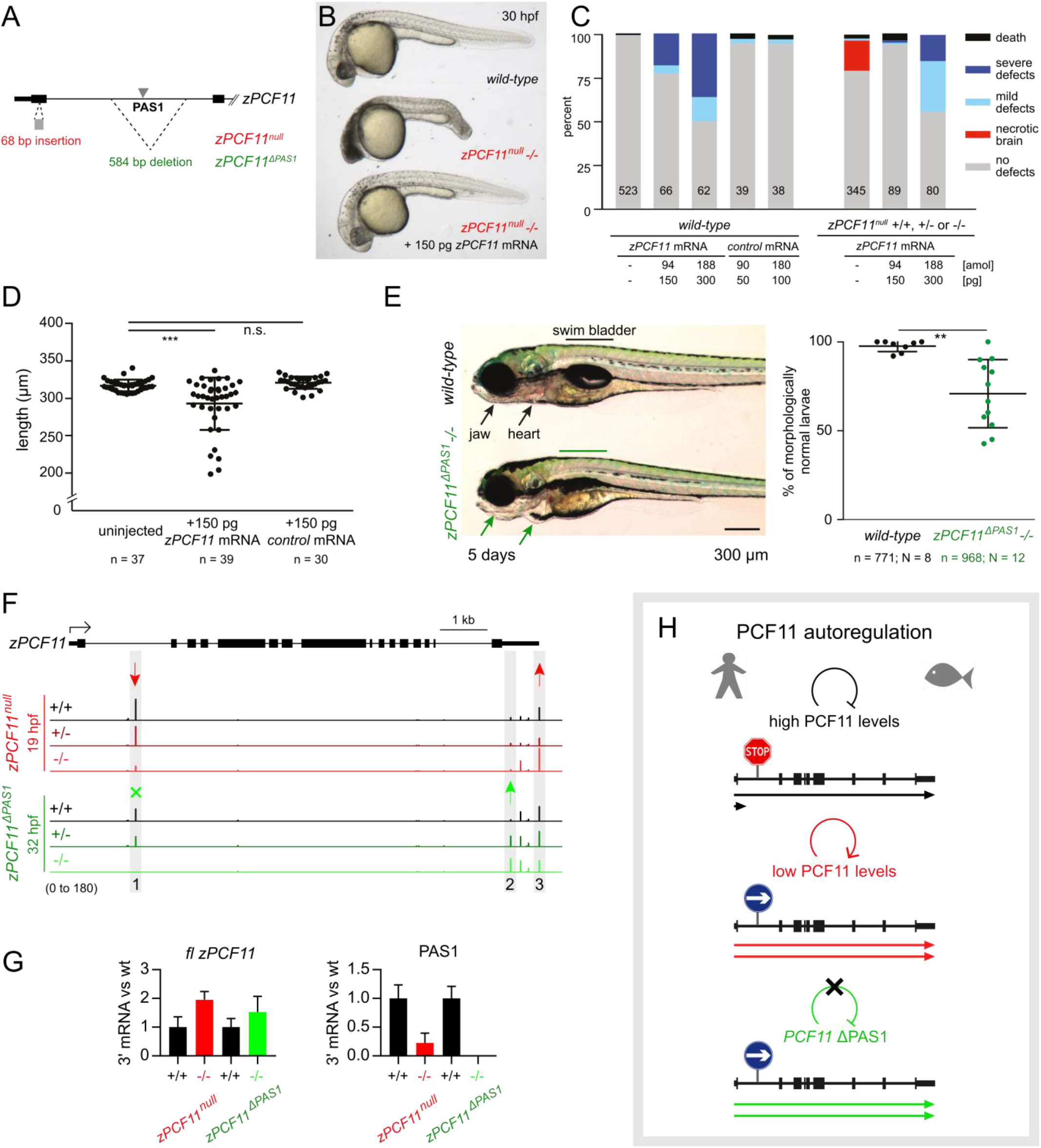
Zebrafish PCF11 is essential for development and undergoes PAS1-dependent autoregulation. (A) Schematic of zebrafish *zPCF11^null^* and *zPCF11^ΔPAS1^* mutants. First two exons of *zPCF11* are shown.
(B) Severe brain necrosis of *zPCF11^null^*−/− embryos is rescued by injection of 150 pg of *zPCF11* mRNA at the 1-cell stage.
(C) Quantification of rescue and overexpression phenotypes upon *zPCF11* mRNA injection. Embryos were scored at 1 day. The numbers within bars indicate the number of embryos scored within each treatment group. (C-D) Control mRNA: *GFP-Bouncer* (Herberg et al., 2018).
(D) Quantification of the decrease in body length at 2 days upon overexpression of *zPCF11* mRNA. Example images of the larvae are shown in Figure S6F.
(E) Phenotypes observed in *zPCF11^ΔPAS1^*−/− larvae at 5 days. (Left) example images; (right) quantification of the morphological defects (lack of swim bladder, heart edema, jaw malformations). n – total number of embryos, N – number of independent crosses. (D-E) Significance determined by unpaired two-tailed t-test.
(F) 3’ mRNA-seq profiles of the *zPCF11* gene for the indicated genotypes (red: siblings derived from *zPCF11^null^*+/− incrosses; green: siblings derived from *zPCF11^ΔPAS1^*+/− incrosses). Average values of 3-6 biological replicates (individual embryo heads, see Figure S6H). Arrows and grey shading indicate significantly altered PAS usage (DEXseq padj < 0.05).
(G) Quantification of *flzPCF11* mRNA (left) and PAS1 usage (right) in indicated mutants relative to the corresponding wild-type. 3’ mRNA-seq was used for quantification (n>=3).
(H) Model of PCF11 autoregulation in human and zebrafish. (Top) When PCF11 protein levels are high (in wild-type situation), *PCF11* transcription is partially attenuated by PAS1 usage and premature termination; as a result only a fraction of transcripts are full-length. (Middle) When PCF11 protein levels are low, PAS1 usage drops leading to more full-length *PCF11* mRNA formation. (Bottom) PAS1 removal leads to increased full-length mRNA and protein production.

Interestingly, our mRNA rescue experiments revealed that injection of higher amounts of *zPCF11* mRNA (300 pg) into embryos derived from *zPCF11^null^*+/− incrosses or overexpression of zPCF11 in wild-type embryos caused a range of morphological abnormalities, including an overall shortening of the body axis (Figure 6C-D, S6F). Thus, both too much and too little zPCF11 interfere with normal development. The necessity of tightly controlled zPCF11 levels *in vivo* prompted us to investigate the importance of the conserved intronic PAS1 in balancing zPCF11 during zebrafish development. To this end, we generated zebrafish mutants, in which the conserved intronic PAS1 was deleted by CRISPR/Cas9 (*zPCF11^ΔPAS1^*) (Figure 6A, S6B-C). Homozygous mutant *zPCF11^ΔPAS1^*−/− larvae showed reduced fitness as evidenced by delayed swim bladder formation, mild jaw abnormalities and weak edema formation at 5 days (Figure 6E). Although the majority of *zPCF11^ΔPAS1^*−/− larvae developed into phenotypically normal adults, the presence of a larval phenotype in the absence of PAS1 suggests that PAS1 might contribute to the PCF11 autoregulatory feedback loop *in vivo*. Consistent with a negative regulation of zPCF11 by PAS1, *zPCF11^ΔPAS1^* −/− embryos showed increased zPCF11 protein levels when compared to wild-type embryos (Figure S6G).

To confirm that PAS1-mediated PCF11 autoregulation occurs during zebrafish development, 3’ mRNA-seq was performed on *zPCF11^null^* and *zPCF11^ΔPAS1^* mutant vs. wild-type embryos. To this end, heads of 3-6 single embryos were sequenced individually for each genotype (+/+, +/− and −/−). 3’ mRNA-seq of *zPCF11^null^*−/− embryos consistently showed that lack of zPCF11 protein leads to 4-5-fold lower PAS1 usage, and a concomitant 2-fold increase in *fl zPCF11* mRNA (Figure 6F-G, S6H). In contrast, and consistent with our immunostainings (Figure S6G), *zPCF11^ΔPAS1^*−/− embryos showed an about 1.5-fold increase in *fl zPCF11* mRNA levels (Figure 6F-G and S6H).

At a global level, *zPCF11^null^* and *zPCF11^ΔPAS1^* mutant zebrafish embryos revealed few statistically significantly changes in APA (Figure S6I). This is likely at least in part due to the fact that zebrafish embryos comprise a heterogenous cell population; analysis of average PAS usage could therefore result in high variability of detected PAS usage between the individual wild-type embryos (Figure S6H, black tracks). Nevertheless, we observed a general tendency for more distal APA in *zPCF11^null^*−/− mutants, and proximal APA in *zPCF11^ΔPAS1^*−/− mutants (Figure S6I), consistent with our findings in human cells (Figures 2B and 5I). The *zPCF11* gene appears to be an APA-prone gene during zebrafish embryogenesis (Figure 6F) as there was a significant distal APA shift in the *zPCF11* 3’UTR in *zPCF11^nul^*−/− embryos, and a proximal shift in *zPCF11^ΔPAS1^*−/− embryos. These data together suggest that zPCF11 favours proximal PAS usage, like its human homolog.

We conclude that both in human cell lines and during zebrafish development, PAS1-linked premature termination promotes PCF11 autoregulation and homeostasis (Figure 6H), which increases animal fitness.

### Transcriptional regulators are controlled by PCF11-dependent premature CPA and termination

We demonstrate above that PCF11 regulates its own expression by premature termination, and also favours proximal PAS usage and early termination genome-wide. This underlines the possibility that PCF11 might also attenuate the transcription of other genes. To validate this hypothesis, we searched our human datasets for protein coding genes significantly upregulated upon PCF11 depletion, which at the same time showed decreased intragenic CPA (PAS usage) and/or decreased intragenic termination (i.e. T4ph mNET-seq signal, Figure 7A). 218 genes were identified as candidates for hPCF11-induced attenuation. 55 genes showed simultaneously decreased CPA and termination (Figure S7A). Premature termination in the absence of a detectable polyadenylated product most likely reflects the unstable nature of such transcripts (Chiu et al., 2018). In the reverse scenario, decreased intragenic PAS usage without associated changes in termination possibly reflects the uncoupling of CPA and termination as described in Figure 2. Strikingly, GO-term analysis of the genes attenuated by PCF11 (Figure 7B) revealed that they are highly enriched for regulators of gene expression, both at the level of transcription and RNA processing. This enrichment was observed independently for both the CPA and termination criteria (Figure S7C-D). We then repeated the CPA-based analysis for the zebrafish dataset and identified 108 putative zPCF11-attenuated genes, which were again enriched for genes involved in transcription (Figure S7E). Notably, PCF11-dependent early CPA can be observed on the very same genes in human and zebrafish, in particular for 3’ mRNA processing factors (Figure 7C-D and S7B). Thus, both in human cells and during zebrafish development, PCF11 downregulates a subset of important transcriptional regulators by premature CPA/termination.

**Figure 7.**
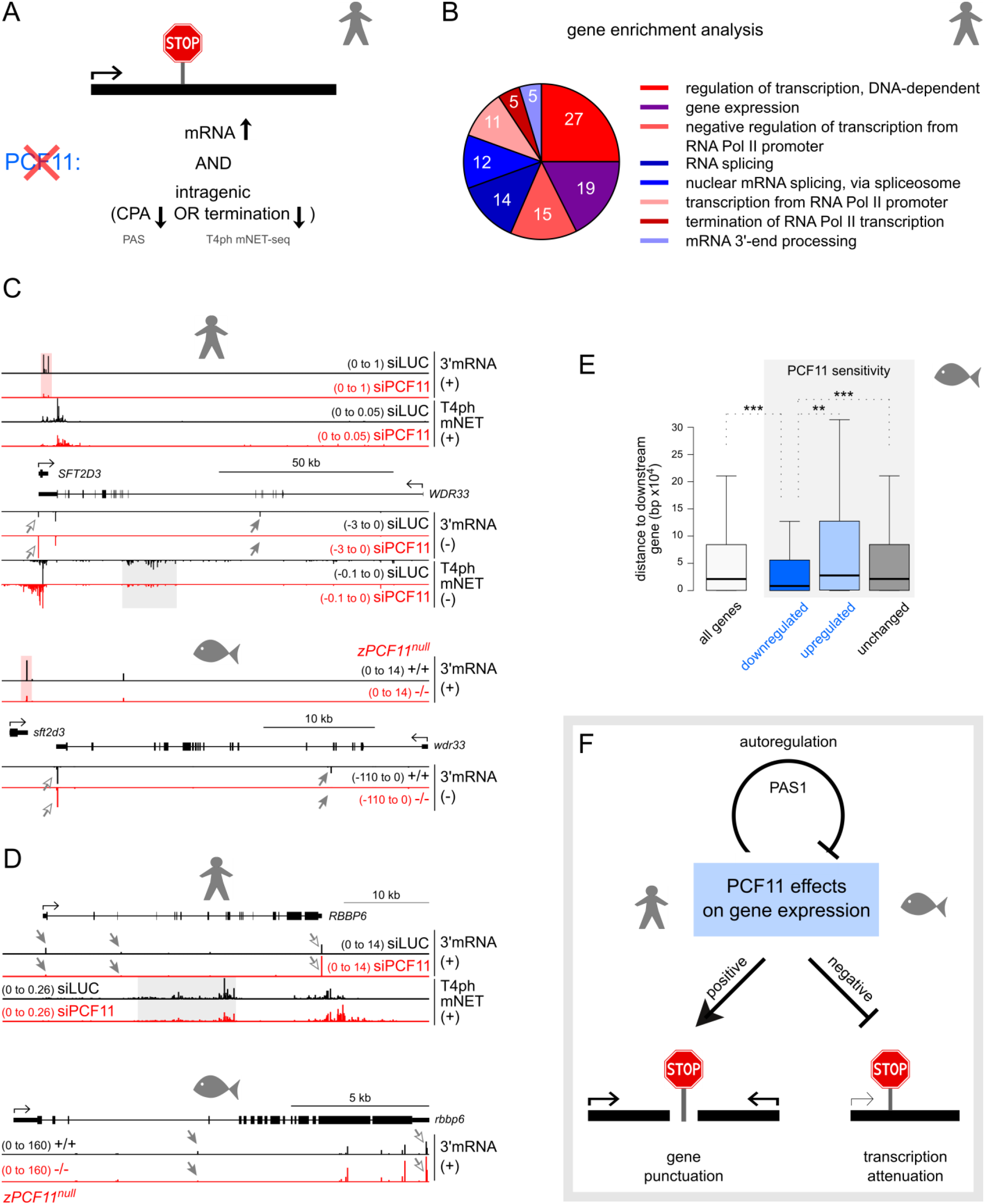
Transcriptional regulators are controlled by PCF11-dependent premature CPA and termination. (A) Criteria for identifying genes attenuated by PCF11-dependent premature CPA/termination: genes which are significantly upregulated upon PCF11 depletion (DEseq padj < 0.05), and harbour either a significantly decreased intragenic PAS (DEXseq padj < 0.05) or show a > 2-fold decreased intragenic T4ph mNET-seq signal.
(B) Enrichment analysis of GO Biological Process for transcripts attenuated by PCF11 in human cells. Numbers in pie chart correspond to number of genes in each category. GeneCodis3 software was used (padj < 0.01 and gene number > 2). Red shades, genes related to transcription; blue shades, genes related to RNA processing.
(C and D) Genomic profiles ± PCF11 of *WDR33* (C) and *RBBBP6* (D) for human cells (top) and zebrafish embryos (*zPCF11^null^*, bottom). Grey shading highlights decreased intragenic T4ph mNET-seq signal and arrows distal APA in PCF11 depleted conditions (grey arrowheads: decreased intragenic PAS signal, white arrowheads: increased 3’ UTR PAS usage).
(E) Analysis of gene distance to downstream gene for genes significantly downregulated, upregulated or unchanged in *zPCF11^null^* −/− vs +/+ embryos. Statistical significance was tested by the Mann-Whitney test.
(F) Model: PCF11 activity displays opposing functions in gene expression. PCF11 punctuates closely spaced genes, leading to their gene expression enhancement. In contrast, PCF11 negatively affects the expression of a subset of transcriptional regulators by attenuating their transcription. PCF11 is also autoregulated by PAS1-dependent premature CPA and termination.

Finally, we sought to determine whether PCF11 positively affects closely-spaced genes in zebrafish, as in human. Indeed, gene distance analysis revealed that zPCF11-sensitive downregulated genes have a significantly closer downstream neighbour, compared to both zPCF11-insensitive and upregulated genes (Figure 7E; note also the downregulation of human and zebrafish *SFT2D3* in Figure 7C).

We conclude that selective functions of PCF11 in punctuating closely-spaced genes, enabling their efficient expression, and attenuating transcription of gene expression regulators are conserved in vertebrates from zebrafish to human (Figure 7F).

## DISCUSSION

### The role of PCF11 in PAS selection and transcriptional termination

This work provides a systematic study of PCF11 in vertebrate gene expression. PCF11 is indispensable for zebrafish development, while in human cells it is required for efficient CPA and transcription termination genome-wide. Furthermore, PCF11 levels in human cells and zebrafish embryos determine alternative polyadenylation patterns: decreased PCF11 levels result in globally more distal PAS usage, whereas increased PCF11 levels (as a result of PAS1 deletion) induced proximal PAS usage. This is in agreement with a previously published mouse screen (Li et al., 2015). We observed fewer significant APA events in zebrafish embryos compared to human cell culture. This may be due to technical reasons (see above), or alternatively APA regulation may reflect an enhanced role for PCF11 in evolution. Accordingly, experiments in *S. pombe* revealed a role for Seb1, but not Pcf11, in the alternative generation of 3’ ends on selected transcripts (Larochelle et al., 2017).

It is notable that PCF11 protein levels are substoichiometric in multiple human tissues and cell lines: an order of magnitude lower than the average number of molecules per cell of other CPA complex subunits. This suggests that human PCF11 is not a core subunit of the CPA complex, but rather an accessory or regulatory factor. In line with this, a previous biochemical study reported that the association between CFIIm and the CPA complex appears weak and transient (Shi et al., 2009).

We suggest that PCF11 acts selectively. Thus, PCF11 ChIP-seq analysis shows binding to a large proportion of transcription units - including transcript classes that are processed independently of the CPA machinery. At the same time, PCF11 binding was undetectable on some active genes undergoing polyadenylation. In our experiments we employed two independent polyclonal antibodies, therefore it is unlikely that our results are due to epitope inaccessibility. However, it is possible that antibody sensitivity, fast RNA processing or transient PCF11 binding preclude PCF11 ChIP detection on some genes. In addition to a selective PCF11 ChIP profile, we found that transcription termination defects and APA changes upon PCF11 depletion were wide-spread, but not universal. Two previous findings support the view that PCF11 in higher eukaryotes is a selective factor. Firstly, staining of *Drosophila* polytene chromosomes with PCF11 antibody correlated with Pol II staining, but fewer bands were visible (Zhang and Gilmour, 2006). Secondly, on the HIV provirus, PCF11 was required for 5’LTR but not 3’LTR termination (Zhang et al., 2007). Therefore PCF11 may act as a selective 3’ processing and termination factor in higher eukaryotes.

If PCF11 acts only on a subset of genes, alternative factors may exist to substitute its functions. Possible candidates are the uncharacterized mammalian CID containing proteins SCAF4/8, RPRD1A/B and/or RPRD2, which are more abundant than PCF11 in human cells and tissues. In *S. pombe* both PCF11 and the SCAF4/8 homolog Seb1 independently contribute to CPA and transcription termination (Lemay et al., 2016; Wittmann et al., 2017).

### Regulation of PCF11 expression

Transcription of the *PCF11* gene is tightly regulated, consistent with the possible regulatory nature of PCF11. *PCF11* contains a strong and evolutionary conserved PAS within its first intron (PAS1) which in both human cell culture and zebrafish embryos enables autoregulation by premature CPA and termination. PAS1 renders *PCF11* expression also sensitive to transcription and RNA processing dynamics. Accordingly, slow Pol II elongation, splicing inhibition, or UV treatment lead to almost exclusive transcription of the *sPCF11* isoform, attenuating *flPCF11* expression (Figure S7F). We also noticed that PCF11 mRNA levels fluctuate widely in control cells (Figure S5B). As *PCF11* mRNA is both non-abundant and unstable, transcriptional changes of *PCF11* might have relatively fast functional effects. We speculate that in conditions where elongation or RNA processing are suboptimal (e.g. after UV damage) it is beneficial to down-regulate PCF11 and so reduce CPA and termination efficiency. This may act to counteract the global shortening of transcripts that occurs under such conditions (Devany et al., 2016; Williamson et al., 2017).

### Premature termination as a regulatory paradigm in vertebrates

PCF11 activity displays opposing functions in gene expression. It positively affects expression levels of many genes - especially closely spaced ones which cannot employ failsafe termination mechanisms. Interestingly, such genes are also prone to defective termination under cellular stress conditions (Vilborg et al., 2017). In contrast, we find that unperturbed cells downregulate a subset of genes by PCF11-mediated transcriptional attenuation, and this function is conserved in vertebrates from zebrafish to human. The phenomenon of negative gene regulation by premature termination is well described in *S. cerevisiae*, where it plays a physiological role in response to changing growth conditions (Bresson et al., 2017). In the mammalian system, genome-wide premature termination of protein coding gene transcription has been previously reported in cells depleted for U1 snRNP (Kaida et al., 2010) and upon DNA damage (Devany et al., 2016; Williamson et al., 2017). Interestingly, two studies using different methodology provided evidence that Pol II accumulation at human and *Drosophila* promoters is not solely due to Pol II pausing, but also associated with premature termination (Nojima et al., 2015; Krebs et al., 2017). Therefore, we predict that premature termination is potentially a wide-spread gene regulatory mechanism in higher eukaryotes. Premature termination of protein coding genes in budding yeast is mediated by the Nrd1–Nab3–Sen1 (NNS) complex, which additionally auto-regulates Nrd1 levels by attenuating *NRD1* gene transcription when Nrd1 levels are high (Arigo et al., 2006). Given the lack of a direct homolog for the NNS complex in higher eukaryotes, we speculate that in vertebrates PCF11 might be a functional counterpart of yeast Nrd1. Interestingly, yeast PCF11 gene is also controlled by NNS-dependent premature termination (Creamer et al., 2011), and cooperates with the NNS complex (Grzechnik et al., 2015). Overall we predict that transcription attenuation is a fundamental gene regulatory mechanism conserved in all eukaryotes.

## ACKNOWLEDGEMENTS

We thank Paweł Grzechnik for critical reading of the manuscript, Bin Tian, Michael Tellier, Justyna Zaborowska, Shona Murphy, David L.V. Bauer, and N.J.P group members for discussions, and the EMBL GeneCore in Heidelberg for sequencing of Pol II ChIP. This work was supported by a Welcome Trust Investigator Award (107928/Z/15/Z), an ERC Advanced Grant (339170) to N.J.P., and a Marie Curie fellowship from EU FP7 (327985) to K.K.-G. A.P. was supported by IMP and Boehringer Ingelheim, as well as the HFSP (CDA00066/2015) and FWF (START project Y 1031-B28).

## STAR METHODS

### KEY RESOURCES TABLE

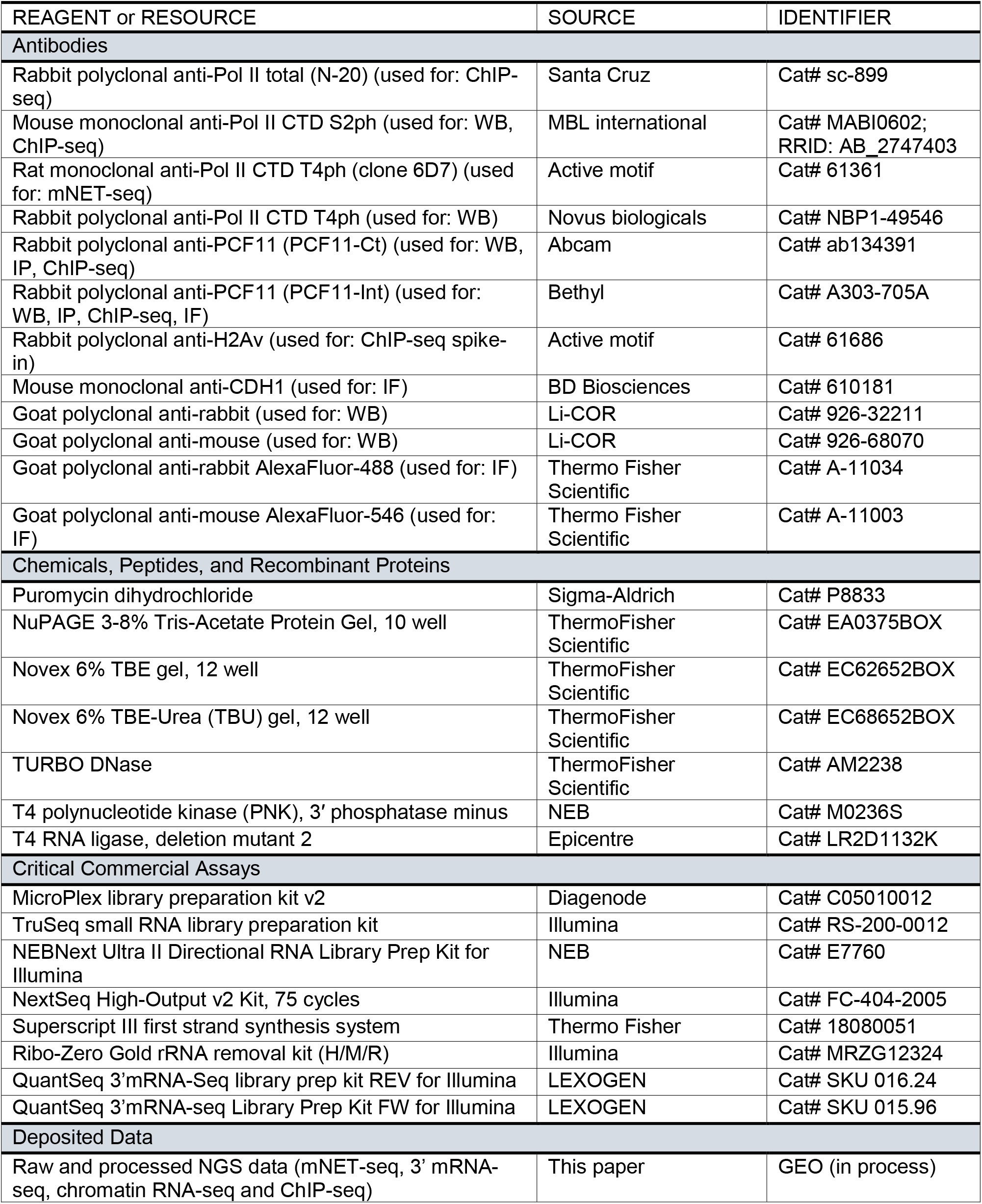
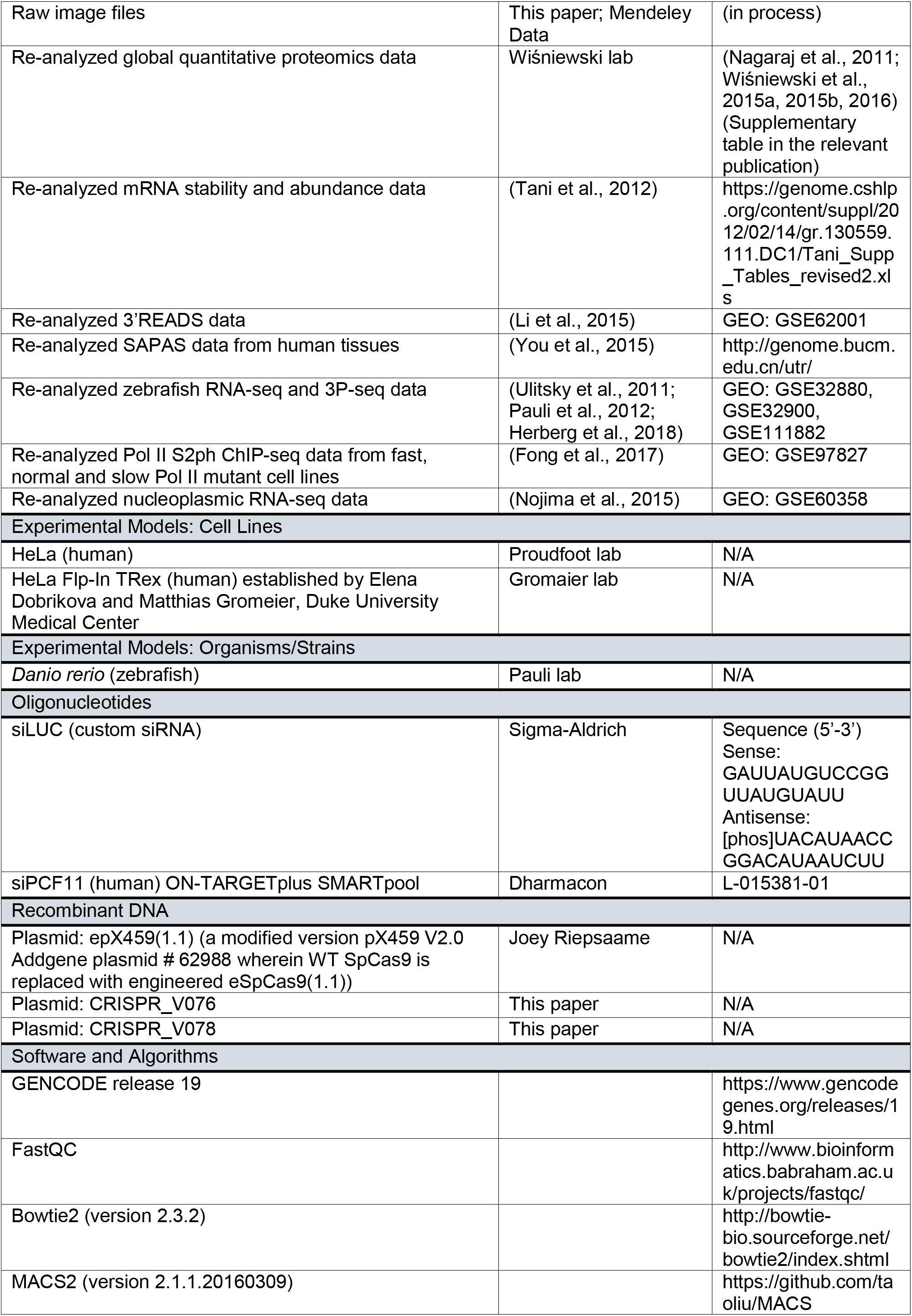
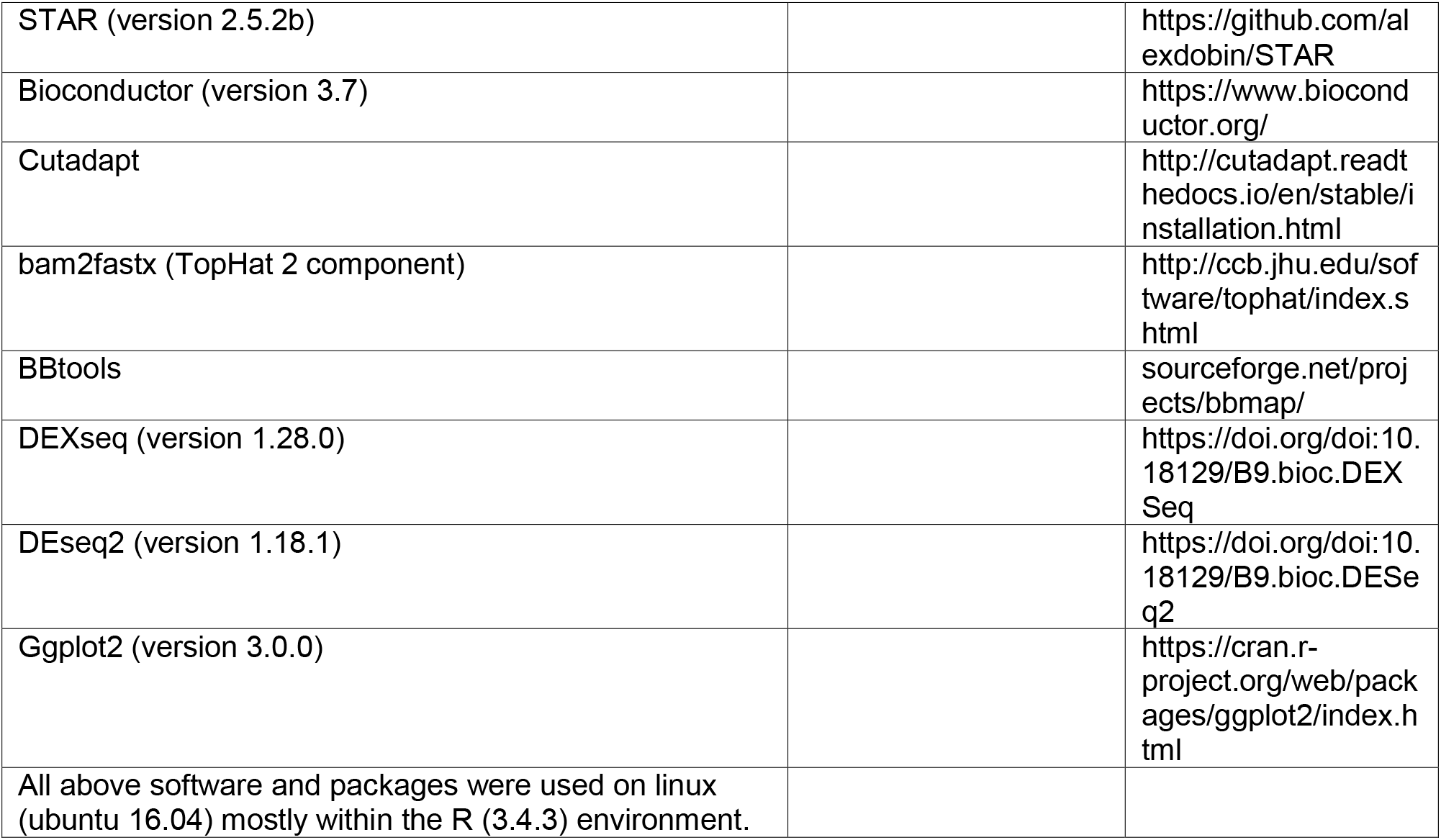

### CONTACT FOR REAGENT AND RESOURCE SHARING

Further information and requests for resources and reagents should be directed to the lead contact, Nicholas Proudfoot.

### EXPERIMENTAL MODEL AND SUBJECT DETAILS

#### Cell lines

All human cell culture experiments were performed in HeLa cells, either wild-type or engineered HeLa Flp-In TRex (established by Elena Dobrikova and Matthias Gromeier, Duke University Medical Center). Cells were cultivated at 37°C and 95% humidity with 5% CO2 in Dulbecco’s Modified Eagle’s Medium (DMEM), high glucose (4,5 g/l) with 10% fetal calf serum (FCS, Perbio) and 1% L-Glutamine (200 mM).

#### Zebrafish

Zebrafish (*Danio rerio*) were raised according to standard protocols (28°C water temperature, 14/10 hr light/dark cycle). TLAB fish, generated by crossing zebrafish AB and the natural variant TL (Tupfel Long-fin) stocks, served as wild-type zebrafish for all experiments. *zPCF11^null^* and *zPCF11*^Δ^*^PAS1^* mutant zebrafish were generated as part of this study and are described in detail below. All fish experiments were conducted according to Austrian and European guidelines for animal research and approved by local Austrian authorities (animal protocol GZ: 342445/2016/12).

### METHOD DETAILS

#### siRNA transfection

siRNA treatment was performed using Lipofectamine RNAimax (Thermo) as described in the product manual. A pool of 4 siRNAs was used to target PCF11 (Dharmacon ON-TARGETplus SMARTpool L-015381-01) as well as a control siRNA against Luciferase (siLUC, see Key Resources Table). The efficiency of depletion was tested by Western blot. Initial time course and a 2-50 nM concentration range test was performed to determine optimal knock-down conditions. All genomic knock-down experiments were performed for 48 hr. 5 nM siRNA concentration was chosen as the standard condition, some replicate genomics experiments were additionally performed using 50nM siRNA, as indicated in genomic dataset description. The 5 nM and 50 nM treatments gave similar genome-wide effects therefore had been treated as biological replicates in the manuscript.

#### Immunoblotting

Proteins were resolved by electrophoresis using 3-8% Tris-Acetate gels (NuPAGE) that separate the migration of PCF11 and Pol II proteins, and blotted onto nitrocellulose membranes. Blots were probed with the antibodies described Key Resources Table, and visualized on a Li-COR Odyssey machine. Li-COR software was used for quantifications.

#### Deletion of *PCF11* PAS1 in human cells by CRISPR/Cas9

Protospacer sequences were cloned into BbsI sites of column-purified plasmid epX459(1.1), a modified version pX459 V2.0 (gift from Feng Zhang (Addgene plasmid # 62988)) wherein WT SpCas9 is replaced with engineered eSpCas9(1.1) (gift from Feng Zhang (Addgene plasmid # 71814)) via KflI/ApaI subcloning. Briefly, equimolar amounts (10 uM; 10 ul) of overlapping oligos harbouring the appropriate sgRNA target sequences were phosphorylated (T4 PNK, NEB) and annealed for 5 min. at 95° before slowly cooling to room temperature. Phosphorylated and annealed oligos were subsequently ligated (T4 ligase, NEB) overnight at room temperature into BbsI-digested epX459(1.1) (5:1 insert-to-plasmid ratio). Upon E.coli (DH10b) transformation and ampicillin selection, plasmid DNA of individual inoculated bacterial clones was prepped (QIAprep Spin Miniprep kit, Qiagen) and correctly cloned protospacer sequences verified using Sanger sequencing (using the primer tandem_sgRNAs_seq – TTCGCCACCTCTGACTTGAGCGT). The following oligos were used for protospacer cloning: PCF11-PAS1_1_F (5’-caccGACCGTCTCTAAACAATATAT – 3’) and R (5’-aaacATATATTGTTTAGAGACGGTC-3’); PCF11-PAS1_2_F (5’-caccGACAAGATACACGGTTTCAGG-3’) and R (5’-aaacCCTGAAACCGTGTATCTTGTC-3’). Guide RNA/Cas9 expression vectors were transfected into HeLa Flp-In TRex cells using Lipofactamine 2000 (Thermo Fisher Scientific) according to manufacturer’s instructions. 24 hr after transfection puromycin was added to the cells at 3μg/ml concentration to select for plasmid-expressing cells. After 24 hr of puromycin selection, the medium was exchanged for non-selective conditions and cells were left to recover for 72 hr before sorting single cells by FACS into four 96 well plates. Individual clones were screened for PAS1 deletion using PCR and the nature of the deletion of candidate clones was verified by Sanger sequencing using below primers (PCF11_PAS1_genotyping_F and R). Initially obtained clones were wild-type, PCF11 PAS1 deletion clones only generated colonies 1-2 weeks after normal clones, indicating possible early cell cycle block in the mutant cells.

PCF11_PAS1_genotyping_F: 5’-TCCCTGATAGCGAAGGAGTG-3’

PCF11_PAS1_genotyping_R: 5’-TGTTGAGTATGACGAATGCTTCC-3’

#### ChIP sequencing

Cells were cultivated on 150 mm dishes until 70% confluency, fixed by addition of 1% formaldehyde for 15 min at 37°C and quenched by addition of glycine (125mM) for 5 min. The cells were collected by scraping on ice, washed 3 times with cold PBS, resuspended in 1.5 ml L1 buffer (50 mM Tris pH 8.0; 2 mM EDTA pH 8.0; 0,1% NP40; 10% glycerol; protease inhibitors) per 10^7^ cells, and lysed on ice for 5 min. The nuclei were collected by centrifugation at 800 g for 5 min at 4°C and lysed in 1,5 ml of L2 buffer (0,2% SDS; 10 mM EDTA; 50 mM Tris pH 8.0; protease inhibitors). The suspension was sonicated in 15 ml conical tubes in a cooled Bioruptor (Diagenode) for 15 min at high settings, and cleared by centrifugation for 10 min at 13000 rpm. The chromatin (DNA) concentration was quantified using NanoDrop (Thermo Scientific) and the sonication efficiency monitored on an agarose gel. Protein A and protein G dynabeads (Thermo Fisher Scientific, combined 1:1) were blocked with BSA (250mg/ml beads) in dilution buffer (0,5% NP40; 200 mM NaCl; 50 mM Tris pH 8.0; protease inhibitors) for 2 hr in cold room. The chromatin was diluted 10x in the dilution buffer. For calibration, 25 ng of Drosophila chromatin was added per 100 μg of human chromatin (Egan et al., 2016). The chromatin was pre-cleared with blocked beads for 1 hr at 4°C. 100 μg of pre-cleared chromatin was incubated with 10 μg of α-PCF11 or α-Pol II antibody and 0.5 μg of Drosophila-specific α-H2Av O/N at 4°C, then with 60 μl blocked beads for further 1-2 hr at 4°C. The beads were washed 2x with WB-150 (0.02% SDS; 0.5% NP40; 2 mM EDTA; 150 mM NaCl; 20 mM Tris pH 8.0), 3x with WB-250 (0.02% SDS; 0.5% NP40; 2 mM EDTA; 250 mM NaCl; 20 mM Tris pH 8.0), 2x with WB-500 (0.02% SDS; 0.5% NP40; 2 mM EDTA; 500 mM NaCl; 20 mM Tris pH 8.0) and finally 1x again with WB-150. The immuno-complexes were eluted by two 15 min incubations at 30°C with 100ul elution buffer (1% SDS, 100mM NaHCO3), and de-crosslinked for 4 hr at 65°C in the presence of 10U RNase A. The immunoprecipitated DNA was then purified with the MinElute PCR purification kit (Qiagen) according to manufacturer’s protocol and used for library preparation. Diagenode MicroPlex library preparation kit v2 (C05010012) was used to prepare libraries for sequencing, following manufacturer’s instructions. Indexed libraries were quantified, normalized and pooled for sequencing on Illumina NextSeq550 system (applies to: PCF11, CPSF73 and Pol II ChIP in wt and deltaPAS1 cells). Pol II ChIP experiments in siLUC and siPCF11 conditions were performed similarly, however without spike-in addition; further on those genomic libraries were prepared using NEBNext ChIP-Seq master-mix kit and sequenced on a 50-bp single-end run using the Illumina HiSeq 2000 platform.

#### Mammalian Native Elongating Transcript sequencing (mNET-seq)

Detailed protocols for mNET-seq and were previously described (Nojima et al., 2015, 2016). In brief, the chromatin fraction was isolated from 3 × 107 HeLa cells. Chromatin was digested in 100 μl of MNase (40 units/ μL) reaction buffer for 5-18 min at 37°C in a thermomixer (1,400 rpm). After addition of 10 μL EGTA (25mM) to inactivate MNase, soluble digested chromatin was collected by 13,000 rpm centrifuge for 5 min. The supernatant was diluted with 400 μL of NET-2 buffer (50 mM Tris-HCl pH 7.4, 150 mM NaCl and 0.05% NP-40) and Pol II antibody-conjugated beads were added. 40 μg of T4ph Pol II antibody was used per sample. Immunoprecipitation was performed at 4°C for 1 hr. The beads were washed with 1 ml of NET-2 buffer six times with 100 μl of 1xPNKT (1xPNK buffer and 0.05 % Triton X-100) buffer once in cold room. Washed beads were incubated in 50 μl PNK reaction mix (1xPNKT, 1 mM ATP and 0.05 U/ml T4 PNK 3’phosphatase minus (NEB) in Thermomixer (1,400 rpm) at 37°C for 6 min. After the reaction beads were washed with 1 ml of NET-2 buffer once and RNA was extracted with Trizol reagent. RNA was suspended in urea Dye (7M Urea, 1xTBE, 0.1% BPB and 0.1% XC) and resolved on 6% TBU gel (Invitrogen) at 200 V for 5 min. In order to size select 30-160 nt RNAs, a gel fragment was cut between BPB and XC dye markers. 0.5 mL tube was prepared with 3-4 small holes made with 25G needle and placed in a 1.5 mL tube. Gel fragments were placed in the layered tube and broken down by centrifugation at 12,000 rpm for 1 min. The small RNAs were eluted from gel using RNA elution buffer (1 M NaOAc and 1 mM EDTA) at 25°C for 1 hr in Thermomixer (900 rpm). Eluted RNA was purified with SpinX column (Coster) with 2 glass filters (Millipore) and the flow-through RNA was ethanol precipitated. mNET-seq libraries were prepared using TruSeq small RNA library preparation kit (Illumina, cat. no. RS-200-0012) and user supplied T4 RNA ligase, deletion mutant 2 (Epicentre, cat. no. LR2D1132K), according to Illumina instructions. 13-15 cycles of PCR were used to amplify the library. Before sequencing, the libraries were size-selected on a 6% TBE gel selecting only the 150-230 bp PCR product to exclude primer-primer ligated DNA. Gel elution was performed as described above. The libraries were sequenced on NextSeq500 using NextSeq High-Output Kit, 75 cycles (Illumina). mNET-seq experiments were performed and sequenced as independent biological repeats: 3 repeats of siLUC and siPCF11 experiments, and 2 repeats for wt and muB PCF11ΔPAS1 cells.

#### Chromatin-bound RNA sequencing (chrRNA-seq)

Chromatin-bound RNA-seq protocol was previously described (Nojima et al., 2015). 1 × 107 cells for each condition were resuspended in 12ml of ice cold PBS. Cells were spun down at 500g, 5 min at 4°C and cell pellets were resuspended in 800μl of HLBN hypotonic buffer (10 mM Tris-HCl pH 7.5, 10 mM NaCl, 2.5 mM MgCl_2_, 0.05% NP40). 480 μl of buffer HLBNS (HLBN, 25% sucrose) was carefully under-layered to create sucrose cushion, and nuclei were isolated by centrifugation for 5 min at 1000g at 4°C. Supernatant containing cytoplasmic debris was discarded and the nuclear pellet was re-suspended in 100 μl of ice-cold buffer NUN1 (20 mM Tris-HCl pH 7.9, 75 mM NaCl, 0.5 mM EDTA, 50% glycerol; 1 mM DTT and cOmplete EDTA free protease inhibitors (Sigma) added fresh). Nuclei were lysed in 1200 μl of ice-cold lysis buffer NUN2 (20 mM HEPES pH7.6, 300 mM NaCl, 7.5 mM MgCl_2_, 0.2 mM EDTA, 1 M urea, 1% NP40; 1 mM DTT) during 15min incubation on ice and RNA-bound chromatin was pelleted at 16000 g for 10min at 4°C. Chromatin-RNA pellet was re-suspended in 200 μl of high salt buffer HSB (10 mM Tris-HCl pH 7.5, 500 mM NaCl, 10 mM MgCl_2_). DNA and proteins were digested with Turbo DNAse (Life Sciences) and proteinase K (10 mg/ml, ThermoFisher, nuclease free), incubating on ThermoMixer at 37°C for 10 min and 30min, respectively. RNA was extracted with 1 ml of TRI Regent (Sigma) according to the manufacturer guidelines. RNA was dissolved in 1xTURBO DNAse buffer, digested with TURBO DNAse for 30 min at 37°C on a ThermoMixer and extracted with TRI reagent. RNA was washed three times with 75% ethanol, and dissolved in water. The RNA integrity was checked on the Agilent 4200 TapeStation system (Agilent Technologies). 1μg of input RNA was depleted of ribosomal RNA with Ribo-Zero Gold Kit (MRZG12324, Illumina) according to manufacturer’s guidelines. 5μl of ribo-depleted RNA (i.e. 12-60 ng RNA according to Qubit quantification) was used as input for library preparation. Chromatin RNA-seq libraries from 2-4 biological repeats were prepared with NEBNext Ultra II Directional RNA Library Prep Kit for Illumina (E7760). Libraries were sequenced on NextSeq500 using NextSeq High-Output Kit, 75 cycles (Illumina). ChrRNA-seq experiments were performed and sequenced as independent biological repeats: 2 repeats of siLUC and siPCF11 experiments, and 4 repeats for wt and muB PCF11ΔPAS1 cells.

#### 3’ mRNA-seq on HeLa cells

PAS mapping (3’ mRNA-seq) was performed on nuclear RNA to enrich for newly transcribed RNAs. To this end, cells on 150 mm dishes were grown until 70% confluent and harvested. After centrifugation, the cell pellet was resuspended in 4 ml of ice-cold HLB+N buffer (10 mM Tris pH 7.5, 10 mM NaCl, 2.5 mM MgCl2, 0.5% NP40) and incubated on ice for 5 min. The suspension was then underlayed with 1 ml of ice-cold HLB+NS buffer (10 mM Tris pH 7.5, 10 mM NaCl, 2.5 mM MgCl2, 0.5% NP40, 10% sucrose) and centrifuged at 420 g for 5 min at 4°C. The supernatant was discarded, and the nuclear pellet washed with PBS. RNA was purified from the nuclei using TRI reagent (Sigma) according to manufacturer’s instructions. Residual DNA was digested using 4U Turbo DNase (Life Tech) for 10 min at 37°C followed by proteinase K digestion for 10 min at 37°C. TRI reagent purification and DNase digestion were repeated. RNA was further acid phenol/chloroform and chloroform extracted, followed by ethanol precipitation. The purified RNA was then resuspended in 20ul ultrapure water. 3’ mRNA-seq libraries were prepared using Lexogen QuantSeq 3′ mRNA-Seq Library Prep Kit REV for Illumina according to manufacturer’s instructions, and sequenced on HiSeq2500. 3’ mRNA-seq experiments were performed and sequenced as independent biological repeats: 4 repeats of siLUC and siPCF11 experiments, and 3 repeats for wt, muA and muB PCF11ΔPAS1 cells. Additionally, one library of further PCF11ΔPAS1 clones muC and muD has been sequenced as well.

#### Generation of *zPCF11^null^ and zPCF11*^Δ^*^PAS1^* mutant fish

*zPCF11^null^* and *zPCF11*^Δ^*^PAS1^* mutant fish were generated by Cas9-mediated mutagenesis. To generate *zPCF11* knockout fish lacking zPCF11 protein, a guide RNA (sgRNA) targeting the first coding exon of the zPCF11 gene was generated according to published protocols by oligo annealing followed by T7 polymerase-driven *in vitro* transcription (gene-specific targeting oligo: zPCF11_ex1_gRNA; common gRNA oligo). To generate zebrafish lacking the conserved PAS1 in intron1 of *zPCF11*, a pool of four sgRNAs targeting intron1 sequences flanking the PAS1 (zPCF11_in1_gRNA1, zPCF11_in1_gRNA2, zPCF11_in1_gRNA3, zPCF11_in1_gRNA4) was generated in a similar manner. SgRNAs were co-injected together with Cas9 protein into the cell of one-cell stage TLAB embryos. Putative founder fish were outcrossed to TLAB wild-type fish. Founder fish carrying germline mutations in the first exon (primer: zPCF11_gt_F1 and zPCF11_gt_R1) or deletions in the first intron (primer: zPCF11_gt_F2 and zPCF11_gt_R2) of *zPCF11* were identified by size differences in the *zPCF11* PCR amplicons in pools of embryo progeny. Embryos from founder fish were raised to adulthood. Sanger sequencing of PCR products of genotyping reactions of adult fin-clips identified the nature of the mutations:

– *zPCF11^null^*: a 68-bp insertion in exon 1, which generates a frameshift mutation after amino acid 28 (H28), and introduces a premature STOP codon after an additional 16 amino acids (MSDDGAREDACREYQSSLEDLTFNSKPH – LVRYQLFQVDNGLSLF*)
– *zPCF11*^Δ^*^PAS1^*: a 584-bp deletion in intron 1, which deletes the entire PAS1.

Homozygous *zPCF11^null^* and *zPCF11*^Δ^*^PAS1^* mutant embryos (*zPCF11^null^−/−* and *zPCF11*Δ*PAS1−/−*

) were generated by incrossing heterozygous adult fish (*zPCF11^null^+/−* or *zPCF11*Δ*PAS1+/−)*. *zPCF11^null^* mutant fish could only be maintained as heterozygotes due to embryonic lethality of *zPCF11^null^−/−* embryos.

#### Genotyping of *zPCF11^null^ and zPCF11*^Δ^*^PAS1^* mutant fish

Genotyping of *zPCF11^null^* fish (68-bp insertion) was performed by PCR amplification of exon 1 of the *zPCF11* gene (primers: zPCF11_gt_F1 and zPCF11_gt_R1). The PCR product size was analyzed by standard gel electrophoresis (wild-type allele: 200 bp, mutant allele: 268 bp).

Genotyping of *zPCF11*^Δ^*^PAS1^* fish (584-bp deletion) was performed by two PCR reactions followed by standard gel electrophoresis. Using PCR reaction 1 (primers: zPCF11_gt_F2 and zPCF11_gt_R2), wild-type fish were reliably identified by the presence of a single 854-bp band. Heterozygous (PCR products of 270 bp and 854 bp) and homozygous (PCR product of 270 bp) fish were, however, not always reliably distinguished as the wild-type allele (upper 854-bp band) in the heterozygous fish was often only very weakly amplified. To identify homozygous fish definitively, PCR reaction 2 was performed using zPCF11_gt_F2 and reverse primer zPCF11_gt_R3, which binds in the intronic region that is deleted in *zPCF11*^Δ*PAS.1*^. This reaction only amplified the WT allele (369 bp), and homozygous mutant fish were therefore easily identified by a complete lack of PCR product.

#### Generation of zPCF11 full-length mRNA

The coding sequence of *zPCF11* was amplified by PCR from cDNA derived from zebrafish embryos (primers: zPCF11_F; zPCF11_R) and cloned by Gibson cloning into the BamHI/EcoRI-digested pCS2+ vector to generate P193: Sp6_zPCF11_SV40-3’UTR. The sequence of *zPCF11* was confirmed by Sanger sequencing. To generate *zPCF11* mRNA, P193 was linearized with NotI, and transcribed using the Sp6 mMessage Machine kit (Ambion). Functionality of the *zPCF11* mRNA was confirmed by the rescue of the fully penetrant brain necrosis phenotype of *zPCF11^null^* −/− embryos by injection of 150 pg into 1-cell stage *zPCF11^null^* −/− embryos.

#### Rescue and overexpression experiment of *zPCF11^null^* mutant zebrafish

*zPCF11^null^* heterozygous incrosses and wild-type embryos were injected with *zPCF11* mRNA (150 pg and 300 pg) and equimolar amounts of control mRNA (*GFP-Bouncer* (Herberg et al., 2018); 50 pg and 100 pg) through the chorion at the one-cell stage. Embryos were scored for morphological defects (e.g. head, tail, and heart defects) and brain necrosis at 1 day post fertilization (dpf) using a stereomicroscope (Zeiss).

For measurement of the length of the body axis, uninjected wild-type larvae and wild-type larvae that had been injected at the 1-cell stage with equimolar amounts of either *zPCF11* (150 pg or 300 pg) or control mRNA (*GFP-Bouncer* (Herberg et al., 2018); 50 pg or 100 pg) were dechorionated at 2 dpf, anesthetized with 0.1% tricaine (E10521, Sigma-Aldrich; 25x stock solution in dH_2_O, buffered to pH 7-7.5 with 1 M Tris pH 9.0) and imaged laterally using a standard stereomicroscope (Zeiss). Body axis length was measured from head to notochord tip using Fiji.

#### Phenotypic scoring of *zPCF11*^Δ^*^PAS1^* mutant zebrafish

*zPCF11*^Δ^*^PAS1^* homozygous mutant fish and wild-type control fish were scored for phenotypic defects and the presence or absence of a swim bladder at 5 days. To this end, larvae were anesthetized in 0.1% tricaine and phenotypically assessed using a standard stereomicroscope (Zeiss).

#### Immunostaining of *zPCF11*^Δ^*^PAS1^* mutant zebrafish

Embryos were fixed at sphere stage in 3.7% PFA at 4°C overnight and washed in PBS-T (0.1% Tween20 in 1x PBS). Before immunostaining, embryos were permeabilized in 0.5% Triton-X-100 in 1x PBS for 1 hr and re-fixed in 3.7% PFA for 20 min with subsequent washings in PBS-T. Embryos were blocked at 4°C overnight (in 20% NGS, 5% DMSO in PBS-T) and stained with a rabbit anti-PCF11 antibody (A303-705A, Bethyl Laboratories, used at 1:40) and a mouse anti-E-Cadherin antibody (610181, BD Biosciences, used at 1:400) at 4°C overnight. Secondary antibody staining was performed at 4°C overnight using goat anti-rabbit AlexaFluor-488 (A-11034, Thermo Fisher Scientific, used at 1:250) and goat anti-mouse AlexaFluor-546 (A-11003, Thermo Fisher Scientific, used at 1:250). DAPI staining was performed for visualize nuclei (incubation with 1x DAPI in PBST for 20 min at room temperature). Embryos were mounted in 1.5% low-melt agarose on a glass-bottom dish (81158, Ibidi) and imaged with an inverted LSM880 Axio Observer confocal microscope (Zeiss), using a 20x objective lens and 1.5x zoom.

#### 3’ mRNA-seq of *zPCF11^null^* and *zPCF11*^Δ^*^PAS1^* mutant zebrafish

Dechorionated embryos of heterozygous *Pcf11*-mutant incrosses were cut in half with a razor blade at 19 hpf (*zPCF11^null^*) or 32 hpf (*zPCF11*^Δ^*^PAS1^*) and, and each head and tail was collected individually in PCR tubes. The anterior halves (heads) were lysed in 10 μl of TCL buffer with 1% beta-mercaptoethanol and flash-frozen on dry ice for subsequent use for RNA isolation and sequencing. The posterior halves (tails) were used for genotyping of each individual sample as described above. Four wild-type, four heterozygous, and six homozygous samples of *zPCF11^null^* and *zPCF11*^Δ^*^PAS1^* mutants were used for library preparation. RNA of selected samples was isolated and purified using Agencourt RNAClean XP magnetic beads (A63987, Beckman Coulter). Strand-specific libraries were generated using the QuantSeq 3’ mRNA Library Prep Kit FW (Lexogen) and used for 100-bp single-end sequencing on the Illumina HiSeq 2500.

### QUANTIFICATION AND STATISTICAL ANALYSIS

#### Human genomic annotation and analyzed gene sets

Hg19/GRCh37 was used as the reference genome. GENCODE release 19 was used for gene annotations: https://www.gencodegenes.org/releases/19.html. This annotation includes 57820 genes (20345 protein-coding, 37475 non-coding). For downstream analysis, we selected a subset of 11947 genes (9095 protein-coding, 2852 non-coding) that satisfied all of the following 3 criteria: 1) had at least one active PAS (see 3’ mRNA-seq analysis below for details); 2) did not overlap with another annotated gene on the same strand; 3) had a 3’ end isolated by at least 6 kb from the downstream annotated gene on the same strand. Those strand-specific isolation criteria allowed to unambiguously assign the directional RNA-seq signal (chrRNA-seq, mNET-seq and 3’ mRNA-seq) to the end of each gene, and also to compute distal APA downstream of annotated gene ends (see below). 6 kb isolation was used because visual inspection of the data in genome browser revealed usage of cryptic non-annotated PASs used upon PCF11 depletion within this window. For meta-profiles and heatmaps, a subset of protein-coding genes longer that 5 kb was used (n=8389), or a further subset of those as indicated in the figure legend. For calculation of distances between genes (Figures 4 and 7) the downstream distance from the gene’s 3’ end to any other annotated gene end (5’ or 3’) on either strand was computed.

#### ChIP-seq mapping, calibration, peak calling and enrichment definition

To allow for detection of proportional changes in global target enrichment between different samples we have added spike-in of *Drosophila melanogaster* chromatin and Drosophila-specific α-H2Av antibody, as described in the experimental methods above. After quality control with FastQC (http://www.bioinformatics.babraham.ac.uk/projects/fastqc/) the curated ChIP-seq reads were mapped with Bowtie2 (http://bowtie-bio.sourceforge.net/bowtie2/index.shtml) using a genome index generated from combined *H. sapiens* hg19 and *D. melanogaster* dm6 genomes. Calculated density plots for distinct samples were normalized to both sequencing depth and the content of Drosophila reads (Egan et al., 2016). MACS2 was used to detect significant enrichments (broad peaks, q-value < 0.01). Peak calling was performed on combined reads from two biological replicates for each antibody. For PCF11, only regions of overlap between the peaks called for the PCF11-Int and PCF11-Ct antibodies separately were considered as PCF11-enriched. PCF11-enriched genes were further defined as a subset of the above described set of 11947 genes (active and separated within the same strand) which gene-body (TSS to PAS) or downstream region (PAS +5kb) overlapped with a PCF11-enriched region. PCF11-enriched genes were considered 3’ enriched if a PCF11-enriched region overlapped the region surrounding the PAS by -2kb to +5kb, independent of possible additional enrichment at the TSS or elsewhere on the gene. All other PCF11-enriched genes were categorized as TSS/gene body enriched.

#### mNET-seq mapping and analysis

Detailed computational mNET-seq workflow has been previously described (Nojima et al., 2016). In brief, reads in FASTQ files were trimmed with Cutadapt using following settings: -a TGGAATTCTCGG -A GATCGTCGGACT -e 0.05 -m 10 --times 1 and mapped with STAR 2.5b to hg19. Last transcribed nucleotides positions from each read were retrieved using in house developed script based on R Bioconductor libraries. Those positions were further used to calculate genome-wide, sequencing depth normalized coverage utilized for downstream analysis and visualization. For premature termination identification, gene bodies of protein-coding genes upregulated upon PCF11 knock-down (DEseq padj < 0.05) were divided into 25 equal bins and T4ph mNET-seq signal was computed in the first 20 bins. In this way we excluded both full-length termination signal downstream of the maximum PAS, as well as the 3’ proximal part of genes, where termination could lead to nearly full-length and possibly functional transcript isoforms.

#### Chromatin RNA-seq mapping and analysis

After quality control with FastQC curated reads were mapped with STAR 2.5b aligner to hg19 (index generated with GRCh37.p13 assembly and gencode.v19.annotation.gtf annotation file). Genomic coverage was normalized to sequencing depth for downstream analysis and visualization.

#### 3’ mRNA-seq mapping

3’ mRNA-seq data were mapped according to the guidelines of QuantSeq library kit manufacturer (Lexogen). Unaligned bam files from HiSeq2500 were converted to FASTQ files with bam2fastx (TopHat 2 component). After overview with FastQC reads were trimmed with BBtools (sourceforge.net/projects/bbmap/) script bbduk using following settings: k=13 ktrim=r useshortkmers=t mink=5 qtrim=r trimq=10 minlength=20. After trimming control with FastQC curated reads were mapped with STAR2.5b aligner to hg19 (index generated with the GRCh37.p13 assembly and gencode.v19.annotation.gtf annotation file).

For PAS calls and APA analysis (below) we have adapted previously published work flows (Derti et al., 2012; Fontes et al., 2017; Rot et al., 2017).

#### Calling polyadenylation sites (PAS)

1. Aligned 3’ mRNA-seq reads were filtered to remove false positives due to internal priming of the QuantSeq assay on genome-encoded poly(A) stretches. To do this, first a crude genomic mask was generated that contained all loci harbouring 6 or more consecutive A bases as well as any 10 nucleotide windows containing more than 6 A bases. For genes expressed from the reverse strand, an analogous T-rich mask was generated. Those crude masks were then corrected to allow for detection of genuine PAS falling in A/T-rich regions by strand-specifically unmasking 20 nucleotide intervals centred at GENCODE 3’ gene ends as well as previously experimentally validated PAS sites detected in all human data sets from (Derti et al., 2012). 3’ mRNA-seq reads falling into those refined strand-specific masks were then removed, and the filtered reads reduced to the most distal nucleotide (3’ end nucleotide).
2. Based on the filtered single-nucleotide 3’ ends strand-specific, sequencing depth corrected, genome-wide density profiles were computed. Density plots from all human samples were summed up (separately for each strand). Because 3’ end cleavage is imprecise, cleavage sites within 30 nt distance were clustered to determine the major active PAS. In detail, first all intervals with at least 30 directly adherent or piled-up filtered read 3’ ends were considered active and merged. The local maximum of such interval was set as the putative PAS. A new 30 nt interval centred around the putative PAS was computed. To avoid overlapping intervals, the merging procedure was repeated, resulting in the final set of 68747 called PAS active in our samples. This set was used for both analysis of alternative polyadenylation (APA) and differential expression analysis.
3. This set of active PAS was then used to count the PAS usage in every sample. The results were normalized for sequencing depth for visualization. APA (DEXseq) and DE analysis were performed on non-normalized reads as the methods used rely on own normalization procedures.

#### Quantification of alternative polyadenylation (APA)

To quantify APA, DEXseq (https://doi.org/doi:10.18129/B9.bioc.DEXSeq) was employed (following previously published workflows for APA analysis (Rot et al., 2017; Fontes et al., 2017). Genes from the analysis set which had at least two alternative active PAS (7718 genes) were subject to differential PAS usage quantification with DEXSeq. Gene coordinates were extended by 6kb downstream of the annotated 3’ end to allow for detection of distal APA beyond annotated gene ends. 4 biological repeats for PCF11 knock-down and 3 biological repeats for PCF11 PAS1 deletion were assayed. Genes where no PAS usage changed significantly between control and treated conditions (DEXseq p-adjusted >= 0.05) were categorized as APA no shift. To determine the direction of APA shift in genes with altered APA, two most statistically differentially used PAS were selected (DEXseq p-adjusted < 0.05), or in the case of genes where only one PAS was significantly altered it was compared to the most frequently used other PAS. If the ratio of the distal to proximal site usage was higher in the treated than in the control cells, the shift was classified as distal, in the opposite situation as proximal (Rot et al., 2017).

#### Analysis of differential gene expression (DE)

Differential expression analysis was performed with DESeq2 (https://doi.org/doi:10.18129/B9.bioc.DESeq2), a well established package from the R Bioconductor project, using the unpaired experimental design. Gene expression was defined as a sum of 3’ mRNA-seq read counts falling into called PAS within above mentioned isolated gene coordinates +6kb. Genes with p-adjusted < 0.05 were classified as differentially expressed.

#### Metagene profiles and heatmaps

Metagene profiles and heatmaps were generated in R, based on GENCODE annotation and enrichment density plots binned into 50 bp intervals, and represent an average of biological replicates. Maximum PAS signal in our datasets was taken as the gene’s 3’ end coordinate. ChIP-seq and mNET-seq data were binned into 50 bp bins. Mean read counts in bins +/− 5kbp from the TSS and -5kb/+10kb from the maximum PAS were extracted for every gene from the above defined analysis gene set limited to protein-coding genes longer than 5 kbp (n=8389). Gene body (GB) of each of gene was divided into 25 equally-sized GB bins and the mean read counts were calculated for each bin. Sequence of bins from genes transcribed from the negative DNA strand was reversed. Finally metagene profiles for distinct genes were assembled by combining respective 5’ end profiles (TSS -5/+1.5 kbp), gene body profiles (GB bins 4:22) and 3’ end profiles (PAS -1.5/+10 kbp). For heatmaps metagene profiles were arranged into matrix sorted based on the signal intensity and visualized with the R ggplot2 package. Mean matagene profiles were calculated as matrices column means.

#### Grouping of genes into proximal and distal major PAS categories

For analysis of association of T4ph mNET-seq signal with proximal and distal APA sites, we selected protein-coding APA genes where the signal of the two strongest PASs differed no more than 2-fold, and that were separated by at least 2 kb, which resulted in a group of 938 genes. The reason for selecting genes with at least two PASs of similar strength was that in case of genes with a very dominant PAS a lack of mNET-seq signal downstream of the minor PASs could be due to a detection limit. The 2kb distance criterion was used, because on average the strongest T4ph mNET-seq signal in control cells is observed within 2kb from the major PAS (see Figure 1C), therefore this distance generally allows to separate T4ph mNET-seq signals from alternative PASs. Those 938 genes were further divided into two groups – one where the major PAS was proximal (PAS1 signal > PAS2 signal, n=413) and second where the major PAS was distal (PAS1 signal <= PAS2 signal, n=525).

#### Zebrafish genomic annotation and 3’ mRNA-seq data analysis

The analyses of zebrafish 3’ mRNA-seq data (read mapping, APA and DE analysis) were all done using the same workflow as established for the human datasets. GRCz10/danRer10 was used as the reference zebrafish genome together with the corresponding ENSEMBL annotation. The read filtering mask used the annotated ENSEMBL 3’ ends. For the *zPCF11^ΔPAS1^* mutant the *zPCF11* gene PAS categorization was manually corrected to APA proximal, as PAS1 was deleted genetically.

#### Data mining and re-analysis of published datasets

Data in Figures S4E-F, S5A and Supplementary table S1 have been extracted from global quantitative proteomics datasets (Nagaraj et al., 2011; Wiśniewski et al., 2015a, 2015b, 2016) from respective supplemental tables.

In Figure 5B, the CPA complex subunits mRNA stability and abundance (as determined by BRICseq) was extracted from table S1 in (Tani et al., 2012).

In Figure S5C, 3’READS dataset (Li et al., 2015) was re-analyzed using the read counts provided by authors in processed table attached to the GEO dataset GSE62001. PAS1 usage was defined as *PCF11* PAS1 reads relative to total *PCF11* 3’READS reads in each sample. This value was then divided by the PAS1 usage of the corresponding siCtrl sample to get the relative PAS1 usage values plotted.

In Figure S5I, PCF11 PAS usage in human tissue was extracted from APASdb (You et al., 2015), which collects datasets specifically profiled for polyadenylation sites using the SAPAS method. Plotted are the processed data as accessed from the database at http://genome.bucm.edu.cn/utr/. For human tissues analysis, tissues were ordered according increasing numbers of PCF11 total PASs mapped i.e. increasing number of transcripts. Colours indicate PCF11 PAS1 usage levels: white/no corresponds to no sequencing counts, yellow/low to 4-10 sequencing counts, orange/medium to 10-30 counts and red/high to >30 counts.

In Figure S6A and E, plotted are zebrafish RNA-seq and 3P-seq data from (Ulitsky et al., 2011; Pauli et al., 2012; Herberg et al., 2018) derived from GEO series GSE32880, GSE32900 and GSE111882.

In Figure 7F, shown are Pol II S2ph ChIP-seq data from (Fong et al., 2017) and nucleoplasmic RNA-seq data from (Nojima et al., 2015). Bigwig files downloaded from GEO series GSE97827 and GSE60358 respectively were directly uploaded to UCSC genome browser to create the genomic profiles.

#### Statistical analysis

Statistical details of experiments are described in the figure legends. n corresponds to the number of genes assayed in a given genomic analysis, or to the number of independent experiments for all other analyses. In all boxplot graphs the bottom and top of the box represent the 25th (Q_1) and 75^th^ (Q_3) percentile respectively and the thick horizontal line the median. The whiskers are defined as:

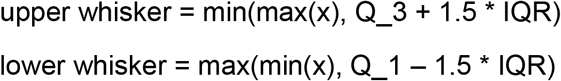

where IQR = Q_3 – Q_1, the box length. Results presented in all bar charts (mainly Western blot quantifications) are expressed as mean values with error bars indicating standard deviation (SD). Statistical significance was analyzed using a Mann-Whitney test or unpaired two-tailed t-test as appropriate, indicated in figure legend. * p <= 0.05, ** p<= 0.01 and *** p<= 0.001 were considered significant.

### DATA AND SOFTWARE AVAILABILITY

The accession number for all NGS datasets (mNET-seq, chromatin RNA-seq, 3’mRNA-seq and ChIP-seq) generated in this paper will be GEO #########. Original images of western blot, gel and immunofluorescent staining assays will be available at Mendeley Data ######. Software used in this work is publicly available under web links indicated in the Key Resources Table.

---

**Figure S1.**
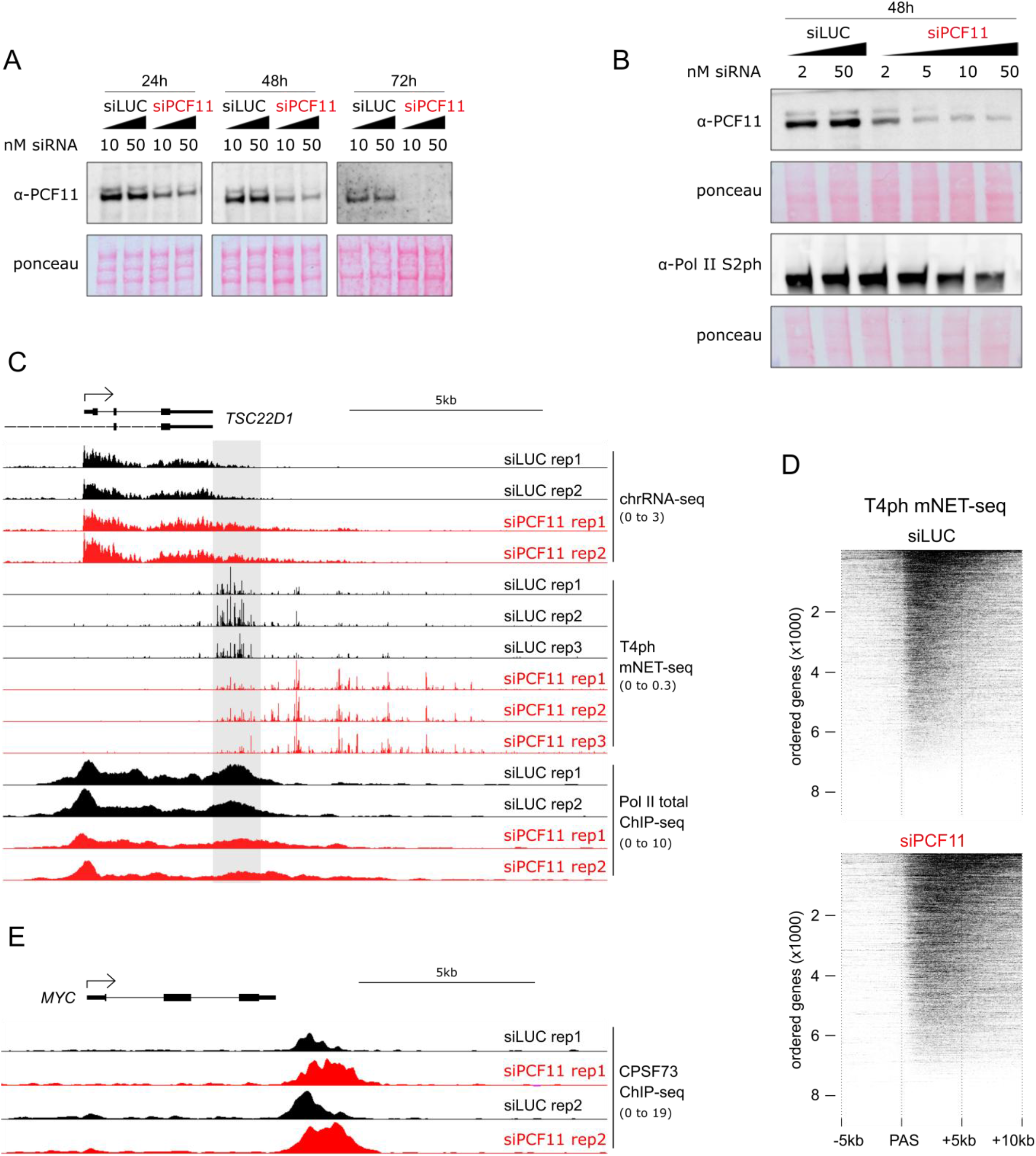
Related to Figure 1. (A and B) Western blot (WB) showing efficiency of PCF11 depletion. Cells were transfected with siRNA against luciferase (siLUC, control cells; black) or siRNA against PCF11 (siPCF11, red). Increasing concentrations of siRNAs are indicated by black triangles. The two WB bands result from PCF11 protein isoforms, as verified by Mass Spectrometry. (A) Time course of siRNA treatment.
(B) Range of siRNA concentrations applied for 48 hr. Note that at high concentration of siPCF11 a decrease in RNA Pol II S2ph signal can be observed.
(C) Genomic profiles showing termination defects in *TSC22D1* upon PCF11 depletion (siPCF11, red) (biological repeats of Figure 1A). For chrRNA-seq and mNET-seq only the sense strand is shown. Grey shading highlights the termination window in control cells (siLUC, black). In all genomic profiles: numbers in brackets indicate the viewing range (rpm).
(D) Heatmaps showing T4ph mNET-seq profiles across individual protein-coding genes ordered based on their T4ph mNET-seq levels (n=8389, gene set as in Figure 1B-D). Top: control cells, bottom: PCF11 depleted cells.
(E) Genomic profile of CPSF73 ChIP-seq binding to *MYC* ±siPCF11.

**Figure S2.**
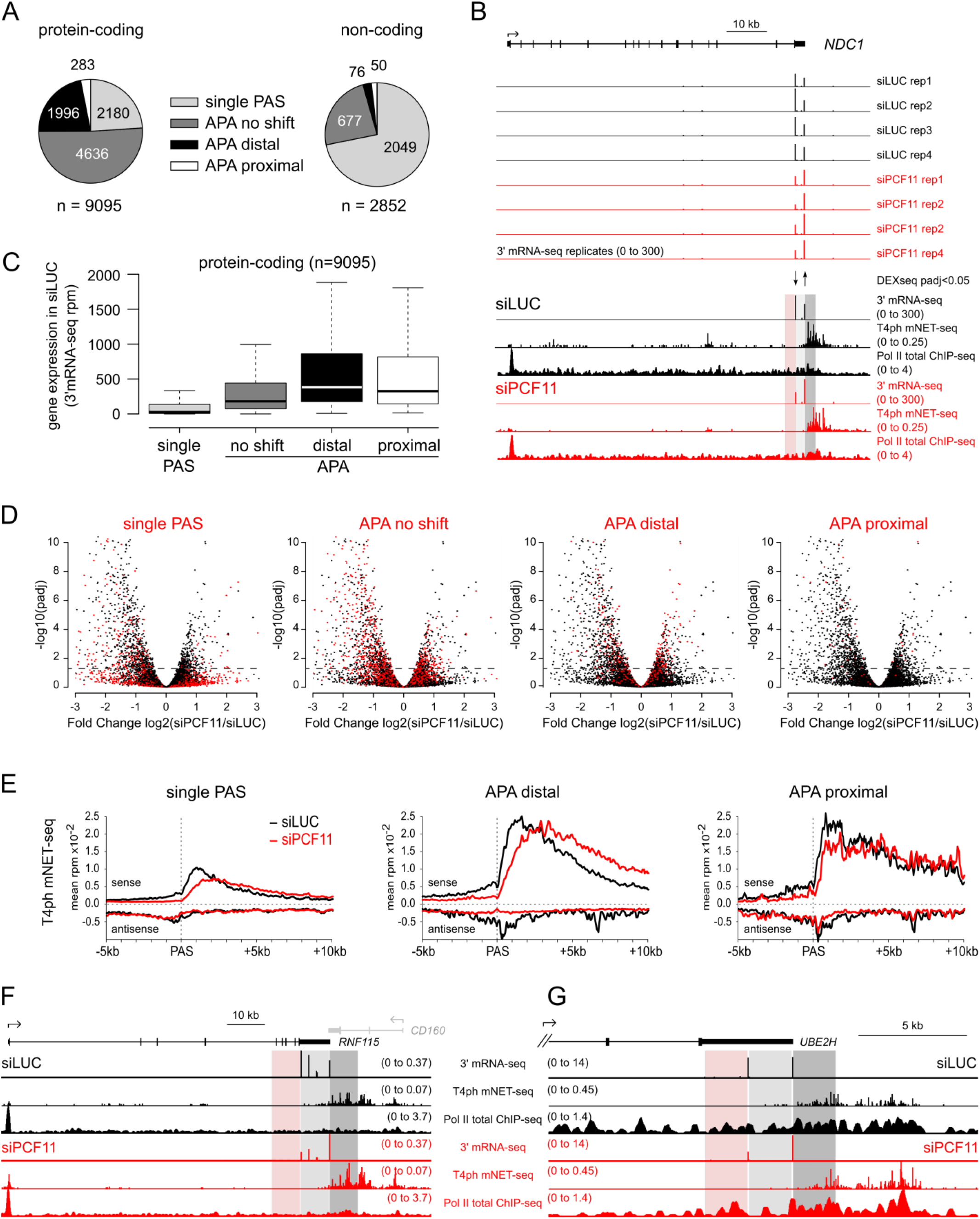
Related to Figure 2. (A) Pie chart of PAS usage upon PCF11 depletion in protein-coding genes (left) and non-coding genes (right): subsets of analysis shown in Figure 2B.
(B) Genomic snapshot of the full *NDC1* locus. Top: biological replicates of 3’mRNA-seq ±siPCF11. Arrows indicate significantly changed PAS usage ±siPCF11 (DEXseq p-adjusted < 0.05). Bottom: average 3’ mRNA-seq, T4ph mNET-seq and Pol II ChIP-seq signal. Red shading highlights the region 2kb upstream of the proximal PAS, light grey the ~2kb region between proximal and distal PASs and dark grey the region 2kb downstream of the distal PAS.
(C) Box plot showing expression levels (measured by 3’ mRNA-seq) of protein-coding genes split into the four PAS usage categories. Here and in all boxplot figures, the thick horizontal line marks to the median, and the upper and lower limits of the box the first and third quartile.
(D) Volcano plots showing differential expression of protein coding genes (n=9095) – negative fold change values: genes down-regulated, positive values: genes up-regulated upon PCF11 depletion. Genes from each PAS usage category (top label) are depicted in red, genes from all other categories shown in black as reference. Genes above the horizontal dashed line have significantly changed expression upon PCF11 depletion (DEseq p-adjusted<0.05 corresponding to -log10(padj)>1.3 in the graph).
(E) Metagene analysis of T4ph mNET-seq signal ±siPCF11 around the major PASs of protein-coding genes in the indicated PAS usage categories. APA no shift category is shown in Figure 2E.
(F-G) Genomic profiles of *RNF115* and *UBE2H*. The shadings highlight the regions: upstream of the proximal PAS (red), in between proximal and distal PASs (light grey) and downstream of the distal PAS (dark grey). The highlighted regions correspond to ~7.5kb for *RNF115* and ~2kb for *UBE2H*.

**Figure S3.**
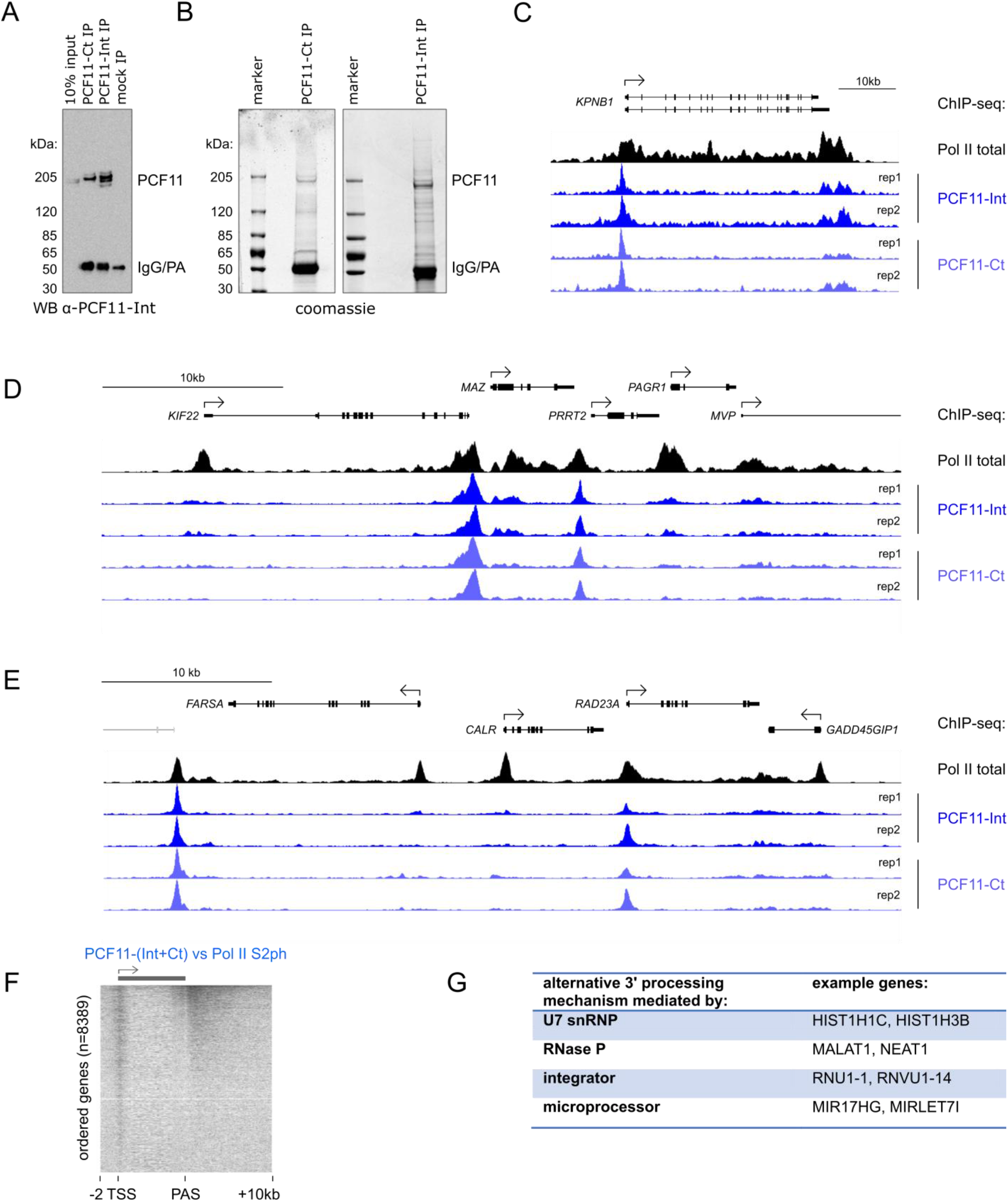
Related to Figure 3. (A) Western blot showing efficiency of PCF11 IP. PCF11 was immunoprecipitated using either α-PCF11-Ct or α-PCF11-Int antibodies. The membrane was probed with α-PCF11-Int.
(B) Coomassie staining of proteins immunoprecipitated with α-PCF11-Ct (left) and α-PCF11-Int antibodies (right). Even under mild conditions (150mM NaCl, 0.5% NP40), the major protein bands just below the 205kDa marker correspond to PCF11 isoforms, as verified by Mass Spectrometry. (A and B) IgG/PA denotes the migration size of immunoglobulin heavy chain (IgG) and protein A (PA).
(C-E) Genomic profiles of PCF11 ChIP-seq binding in the *KPNB1, MAZ*, and *CALR* loci showing 2 biological ChIP replicates using both antibodies (4 ChIP-seq samples total). ChIP-seq for Pol II is shown as reference. Viewing range was autoscaled to maximum signal. Grey outline in (E) corresponds to the inactive *SYCE2* gene.
(F) Heatmap of PCF11-(Int+Ct) ChIP-seq signal normalized to Pol II S2ph across protein-coding genes ranked from highest to lowest PCF11 signal.
(G) Table of alternative 3’ processing mechanism used by non-canonical (non-CPA-dependent) PCF11 target genes.

**Figure S4.**
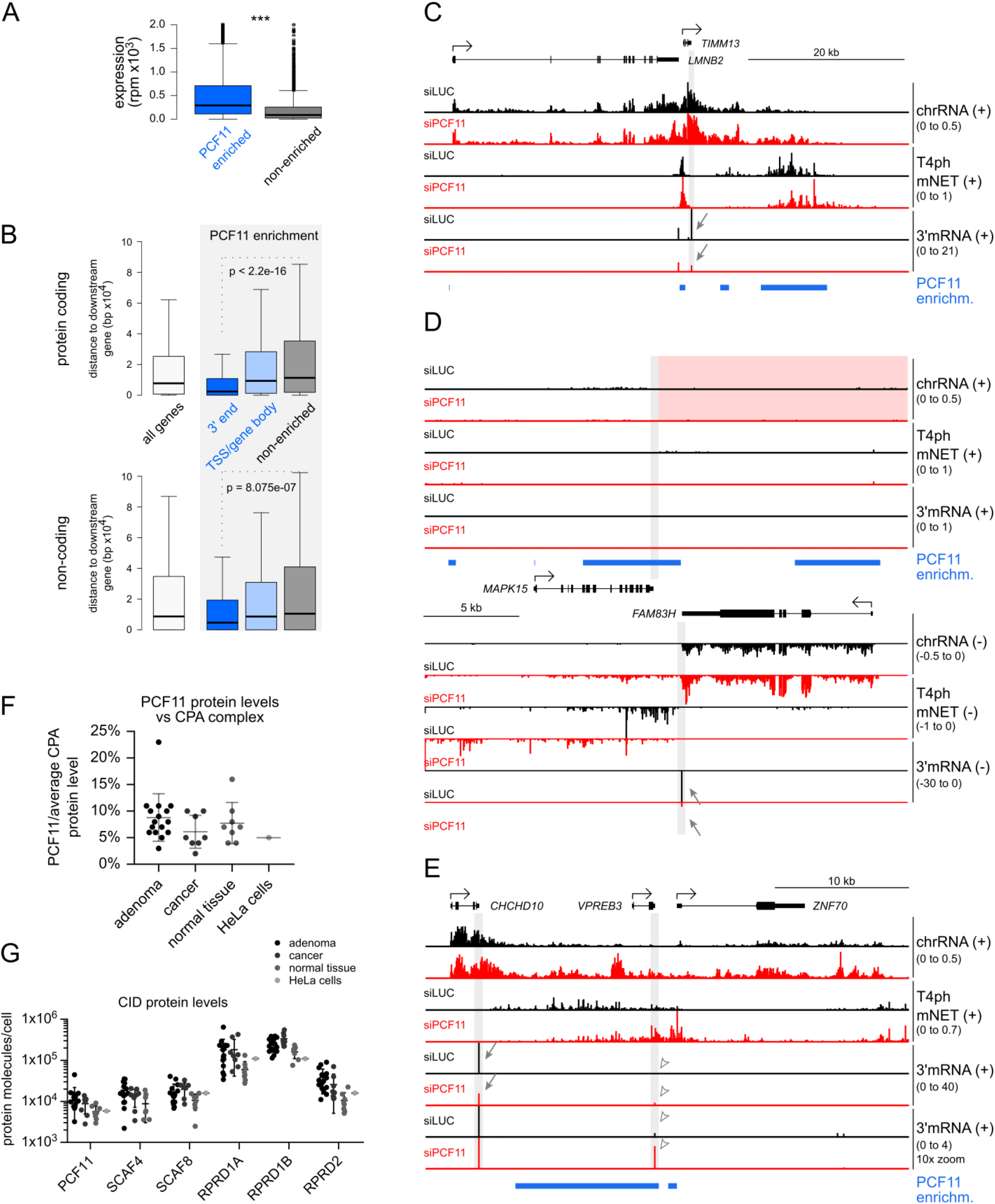
Related to Figure 4 and 5. (A) Comparison of gene expression levels of PCF11 enriched and non-enriched genes based on 3’ mRNA-seq reads.
(B) Boxplots showing the distances between genes’ PASs to their nearest gene downstream. Top: protein coding genes, bottom: non-coding genes. (A and B) Statistical significance was determined using Mann-Whitney test.
(C-E) Genomic profiles of the indicated loci. Grey shading highlights the 3’ ends of relevant genes. Red shading highlights the lack of detectable read-through from *MAPK15* into the highly downregulated *FAM83H*. Blue bars: PCF11 enriched regions; arrows: gene downregulation measured by 3’ mRNA-seq; empty arrowheads: upregulation of *VPREB3* due to read-through transcription from *CHCHD10*.
(F) Quantification of PCF11 protein molecules per cell relative to the average copy number of other CPA complex subunits in biopsies from colorectal adenoma (n=16), cancer (n=8) and normal tissue (n=8), as well as in HeLa cell culture (n=1) based on global quantitative proteomics. PCF11 levels are on average 10-20 fold lower (5-9%) compared to other CPA complex subunits.
(G) Scatter plot of copy numbers of human CID-containing proteins in the same samples as (F). SCAF4 hasn’t been assayed in HeLa cells.
(F and G) Data from (Nagaraj et al., 2011; Wiśniewski et al., 2015). Horizontal lines indicate the mean and SD.

**Figure S5.**
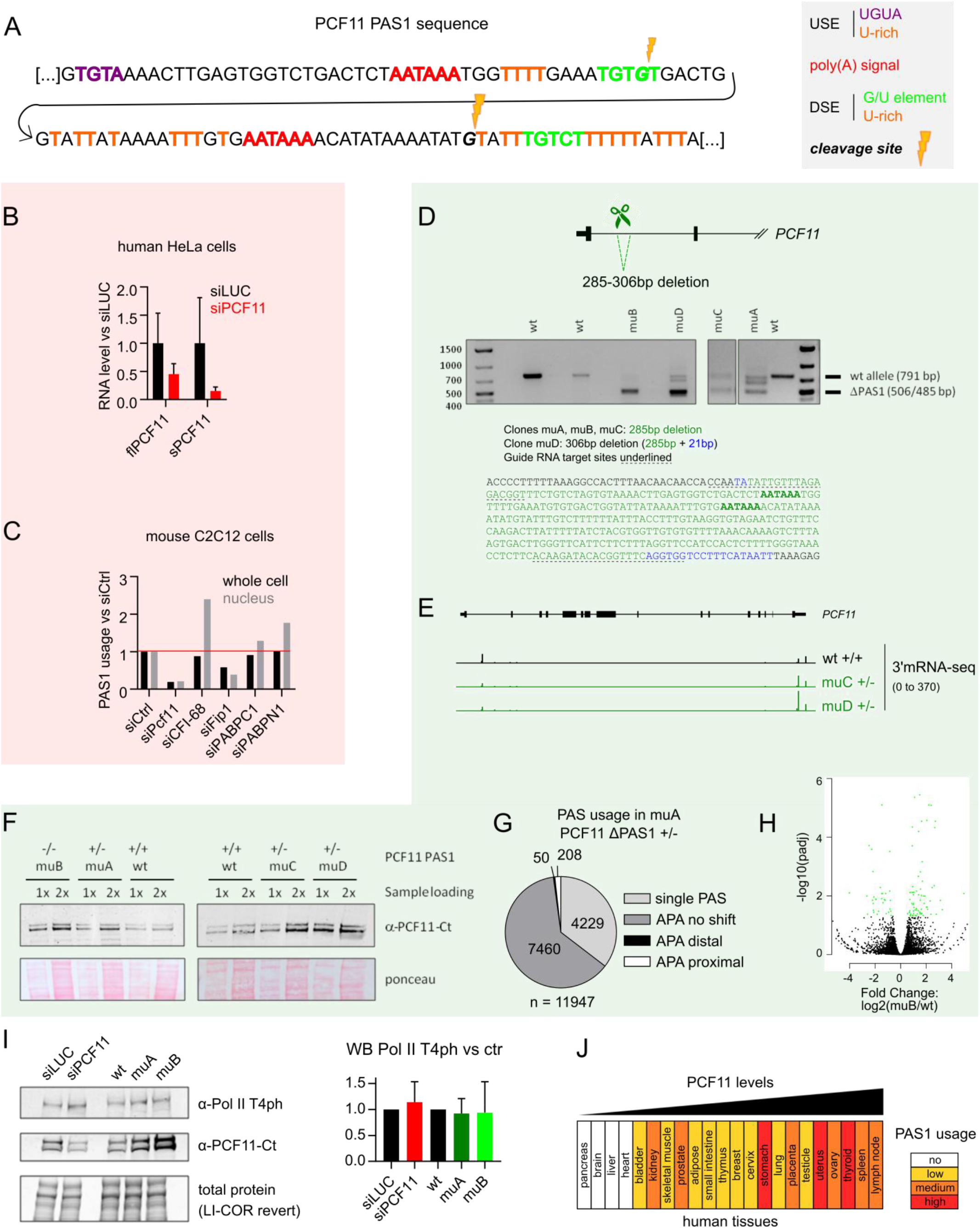
Related to Figure 5. (A) DNA sequence of *PCF11* PAS1. Indicated are the two tandem AATAAA hexamers (red), as well as additional upstream and downstream elements (USE and DSE) forming the poly(A) signal. Two alternative cleavage sites are indicated in *italics* and by the yellow lightning bolt. The second cleavage site is more frequently used.
(B) RNA levels of full-length *PCF11* mRNA (*flPCF11*) and the short *PCF11* isoform (*sPCF11*) in human HeLa cells ±siPCF11 based on 3’mRNA-seq (error bars correspond to SD, n=4). Plotted are values relative to the average levels in siLUC.
(C) *PCF11* PAS1 usage in mouse C2C12 cells depleted of the indicated CPA factors, relative to control cells. Values are based on PAS sequencing using 3’READS method from (Li et al., 2015). Sequenced RNA was extracted either from whole cells (black) or isolated nuclei (grey).
(D) *PCF11* PAS1 deletion using CRISPR/Cas9. (Top) schematic of the deletion, (middle) PCR analysis of the mutant clones showing the degree of PAS1 deletion. Sizes of the wt allele and sequenced deletion alleles are indicated. Only clone muB shows a complete deletion of PAS1. Clones muA, muC and muD are heterozygous, with muD showing the highest proportion of deleted alleles. (Bottom) nucleotides deleted (green and blue) in the four mutant clones based on Sanger sequencing of the PCR products. Underlined nucleotides are CRISPR/Cas9 guide RNA targets, the two tandem poly(A) hexamer signals indicated in bold.
(E) Profile of 3’mRNA-seq on the *PCF11* gene in wt cells and clones muC and muD.
(F) WB of PCF11 protein levels in wt cells and *PCF11* PAS1 mutant clones muA-muD. Each sample was loaded at two concentrations, as indicated.
(G) Pie chart of genome-wide PAS usage and APA occurrence in *PCF11^ΔPAS1^* clone muA vs wt cells (compare with Figures 5I and 2B).
(H) Volcano plot showing differential expression between wt and *PCF11^ΔPAS1^* clone muB cells. Genes indicated by green dots have significantly changed expression upon PCF11 depletion (DEseq p-adjusted<0.05). 2 times more genes are significantly upregulated than downregulated in *PCF11^ΔPAS1^* clone muB vs wt cells.
(I) WB analysis of Pol II T4ph levels in cells where PCF11 is downregulated (siPCF11) or upregulated (PCF11^ΔPAS1^: muA and muB). Left: representative WB, right: quantification of WB experiments, average values relative to control, error bars correspond to SD (n=6 for PCF11 depletion; n=4 for *PCF11^ΔPAS1^*)
(J) *PCF11* PAS1 usage in 22 human tissues. The tissues were ranked according to increasing *PCF11* mRNA levels. Colours indicate *PCF11* PAS1 usage levels: white, no sequencing counts; yellow, 4-10 sequencing counts; orange, 10-30 counts and red, >30 counts. Data extracted from APASdb (You et al., 2015).

**Figure S6.**
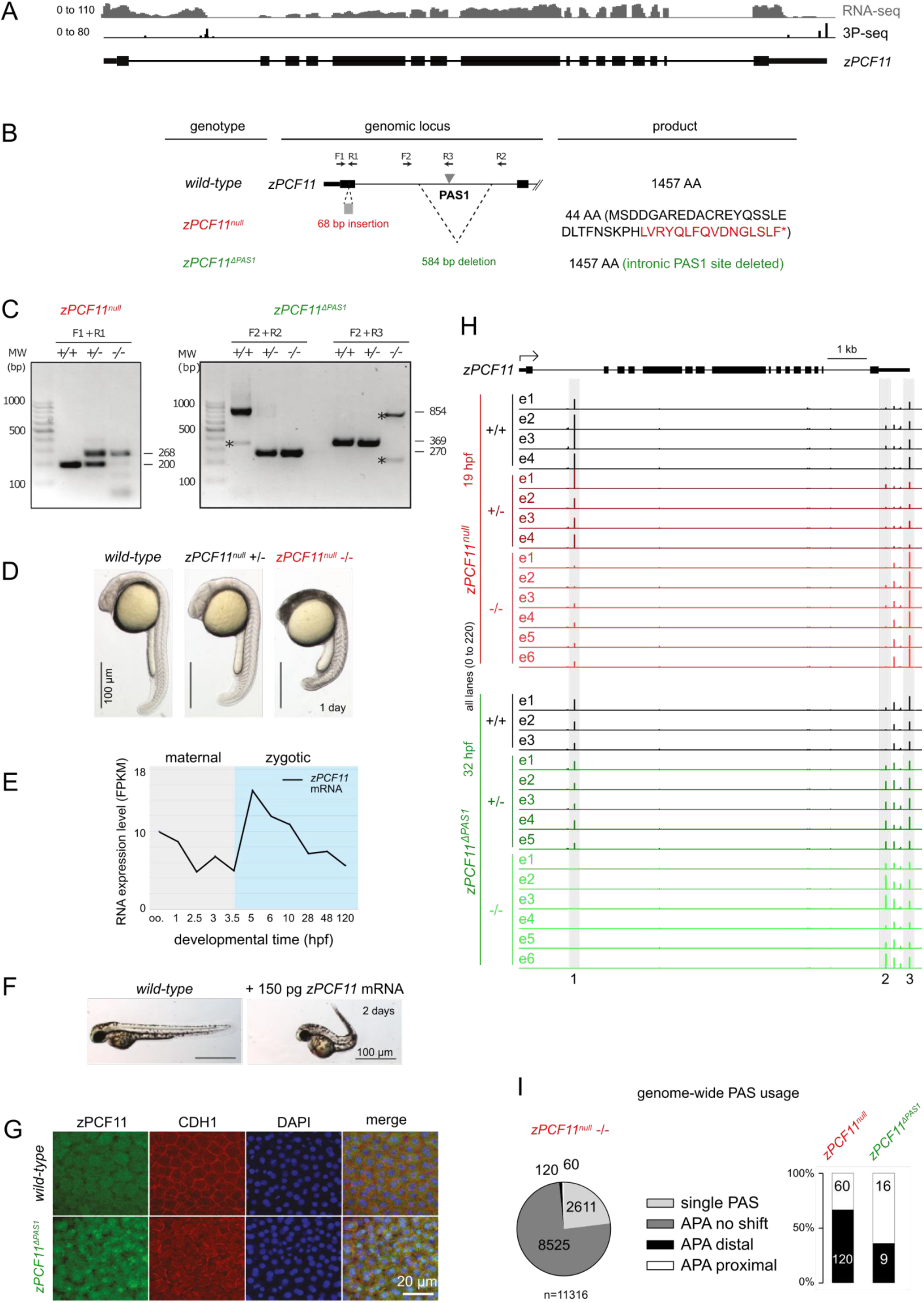
Related to Figure 6. (A) Genomic locus of the zebrafish z*PCF11* gene. Shown is signal of RNA-seq from ovary (GSE111882, Herberg et al., 2018) and 3P-seq from adult fish (GSE32880, Ulitsky et al., 2011).
(B) Overview of zebrafish *zPCF11^null^* and *zPCF11^ΔPAS1^* mutants. The positions of primer sequences used for genotyping and predicted protein products of the wild-type and mutant forms are indicated. Only the first two exons of *zPCF11* are depicted.
(C) Genotyping PCR of wild-type, *zPCF11^null^* and *zPCF11^ΔPAS1^* mutant embryos. Primer locations are shown in Figure S6B. The bands indicated by asterisks are background bands.
(D) *zPCF11^null^−/−* mutant larvae show severe brain necrosis at 1 day.
(E) *zPCF11* mRNA is maternally provided and zygotically expressed. Data from GSE32900 (Pauli et al., 2012) and GSE111882 (Herberg et al., 2018), oo. = oocyte.
(F) Example images of wild-type or *zPCF11* overexpressing larvae at 2 days.
(G) Immunofluorescence of wild-type and *zPCF11^ΔPAS1^*−/− embryos at 5 hours post fertilization. Note the increased PCF11 signal in *zPCF11*^Δ*PAS1*^−/− embryos.
(H) Profile of 3’ mRNA-seq reads from individual zebrafish embryo heads (e1-e6) of the indicated genotypes at the *zPCF11* locus. Grey shadings indicate PASs undergoing APA.
(I) (Left) PAS usage in *zPCF11^null^*−/− mutants vs wild-type. Categories identical to Figure 2A-B. (Right) Significant APA changes in *zPCF11^null^*−/− and *zPCF11^ΔPAS1^*−/− mutants (DEXseq padj < 0.05).

**Figure S7.**
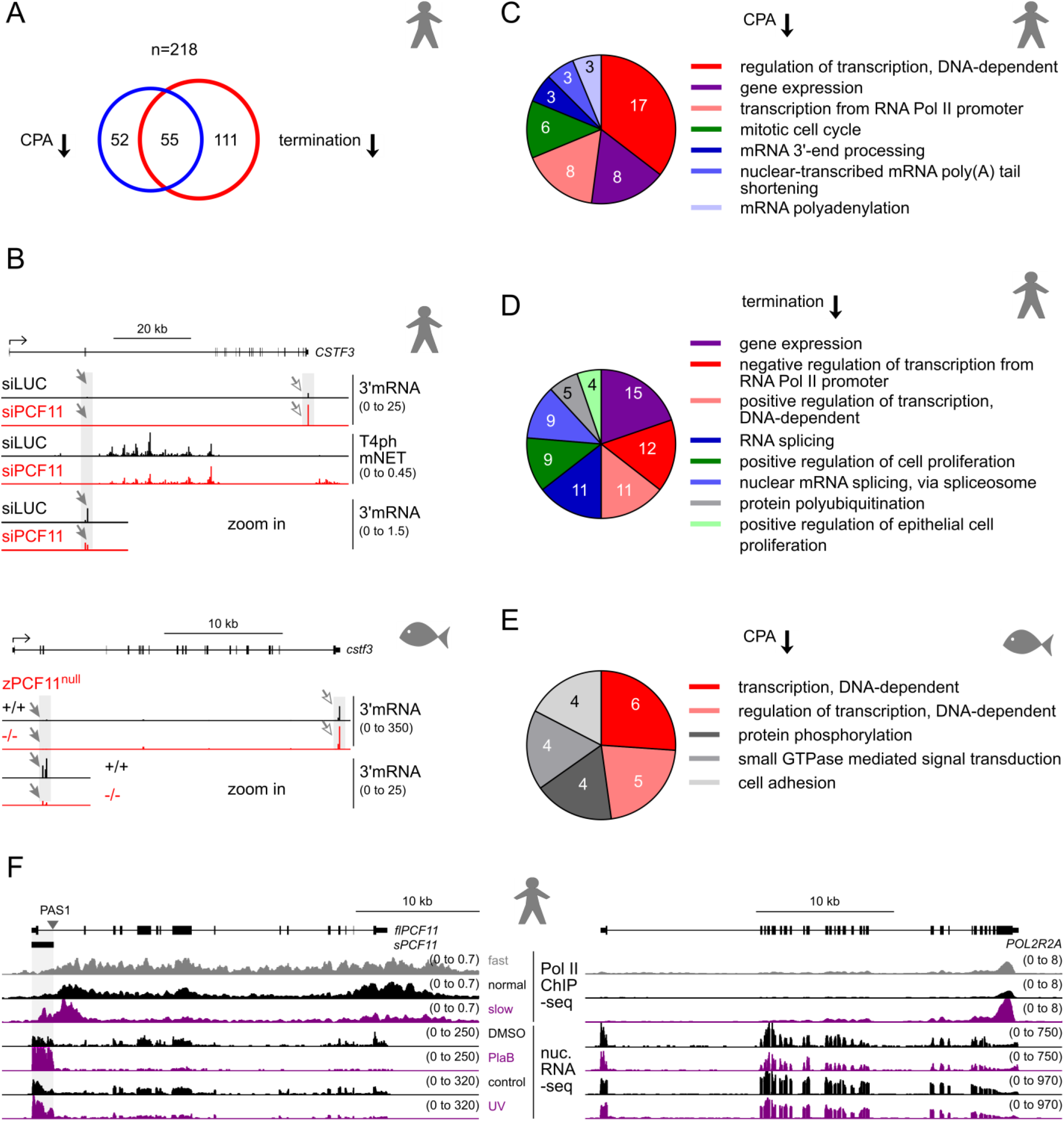
Related to Figure 7 and Discussion. (A) Number of genes and overlap between genes undergoing premature CPA (left) and premature termination (right) in human cells.
(B) Genomic profiles of *CSTF3* gene ±PCF11 for human cells (top) and zebrafish embryos (bottom). Grey shading and arrows highlight distal APA in PCF11 depleted conditions (grey arrowheads: decreased intragenic PAS usage, white arrowheads: increased 3’ UTR PAS usage).
(C to E) Enrichment analysis of GO Biological Process for genes undergoing premature CPA (C) and premature termination (D) in human cells, and genes undergoing premature CPA in zebrafish (E). Parameters used as in Figure 7B. Red shades; genes related to transcription; blue shades, genes related to RNA processing.
(F) Genomic profiles of the *PCF11* (left) and *POLR2A* (right) loci showing the effect of various treatments on the transcription of the two genes. Top: Pol II S2ph ChIP-seq in cells expressing kinetic mutants of Pol II: fast (E1126G), normal (wild-type) and slow (R749H). Data from (Fong et al., 2017). Bottom: RNA-seq of the nucleoplasmic fraction in control cells, cells treated with pladienolide B (PlaB, splicing inhibitor) for 4 hours (Nojima et al., 2015), or cells harvested 2 hours after UV treatment with 20mJ/cm^2^ (UV, T. Nojima). Grey shading highlights the coordinates of the *sPCF11* isoform ending with PAS1. Note that the slow Pol II mutant, splicing inhibition, and UV treatment (purple lanes) all lead to preferential *sPCF11* transcription and concomitant *flPCF11* downregulation.

